# Neutrophil extracellular traps have auto-catabolic activity and produce mononucleosome-associated circulating DNA

**DOI:** 10.1101/2022.09.01.506266

**Authors:** Ekaterina Pisareva, Lucia Mihalovičová, Brice Pastor, Andrei Kudriavstev, Alexia Mirandola, Thibault Mazard, Stephanie Badiou, Ulrich Maus, Lena Ostermann, Julia Weinmann-Menke, Elmo W. I. Neuberger, Perikles Simon, Alain R. Thierry

## Abstract

**Background:** Because circulating DNA (cirDNA) are mainly detected as mononucleosome-associated circulating DNA (mono-N cirDNA) in blood apoptosis has until now been considered as the main source of cirDNA. The mechanism of cirDNA release into the circulation, however, is still not fully understood. This work addresses that knowledge gap, working from the postulate that neutrophil extracellular traps (NET) may be a source of cirDNA, and by investigating whether NET may directly produce mono-N cirDNA

**Methods:** We used the synergistic analytical information provided by specifically quantifying DNA by qPCR, and analyzing fragment size analysis by shallow WGS, and capillary electrophoresis to unequivocally study the following: the *in vitro* kinetics of cell derived genomic high molecular weight (gHMW) DNA degradation in serum; the production of extracellular DNA and NET markers such as neutrophil elastase (NE) and myeloperoxidase (MPO) by *ex vivo* activated neutrophils; *in vitro* NET degradation in serum. We also performed an *in vivo* study in knockout mice, and an *in vitro* study of gHMW DNA degradation, to elucidate the role of NE and MPO in effecting DNA degradation and fragmentation. We then compared the NET associated markers and fragmentation size profiles of cirDNA in plasma obtained from patients with inflammatory diseases found to be associated with NET formation and high levels of cirDNA (COVID-19, N= 28; systemic lupus erythematosus, N= 10; metastatic colorectal cancer, N= 10; and from healthy individuals, N= 114).

**Results:** Our studies reveal that: gHMW DNA degradation in serum results in the accumulation of mono-N DNA (81.3% of the remaining DNA following 24H incubation in serum corresponded to mono-N DNA); “ex vivo” NET formation, as demonstrated by a concurrent 5-, 5- and 35-fold increase of NE, MPO, and cell-free DNA (cfDNA) concentration in PMA-activated neutrophil culture supernatant, leads to the release of high molecular weight DNA that degrades down to mono-N in serum; NET mainly in the form of gHMW DNA generate mono-N cirDNA (2% and 41% of the remaining DNA after 2 hours in serum corresponded to 1-10 kbp fragments and mono-N, respectively) independent of any cellular process when degraded in serum; NE and MPO may contribute synergistically to NET autocatabolism, resulting in a 25-fold decrease in total DNA concentration and a DNA fragment size profile similar to that observed from cirDNA following 8h incubation with both NE and MPO; the cirDNA size profile of NE KO mice significantly differed from that of the WT, suggesting NE involvement in DNA degradation; and a significant increase in the levels of NE, MPO and cirDNA was detected in plasma samples from lupus, COVID-19 and mCRC, showing a high correlation with these inflammatory diseases, while no correlation of NE and MPO with cirDNA was found in HI.

**Conclusions:** Our work thus describes the mechanisms by which NET and cirDNA are linked, by demonstrating that NET are a major source of mono-N cirDNA independent of apoptosis, and thus establishing a new paradigm of the mechanisms of cirDNA release in normal and pathological conditions, as well as demonstrating a link between immune response and cirDNA.

## Background

CirDNA [1] has gained considerable attention in recent decades because of its huge potential for clinical diagnostics. It is already being used for prenatal diagnosis, liquid biopsy in oncology, and other medical applications such as transplantation or non-invasive prenatal testing [2,3]. Further optimization of cirDNA diagnostics would open up significant perspectives for personalized medicine, but that requires an accurate understanding of the fundamental mechanisms underlying the release of DNA into the circulation, of their fragmentation, and of their cellular/tissue of origin.

It was initially thought that cirDNA could be up to 40 kb in size, but was principally 180 bp (or multiples thereof), which corresponds to the size of DNA packed in a mono-nucleosome [4,5]. Observations of mono- and oligo-nucleosomes led to the view that the major mechanism of cirDNA release is apoptosis [3,5,6]. Alternatively, necrosis was reported as playing a major role in response to ionizing radiation [7]. It has also been shown that DNases are the main actors of DNA degradation [8], and the particular endonucleases to which that degradation has been attributed include caspase-activated DNAse (CAD), DNase 1L3, Endonuclease G (Endo G), and DNase I [9–14]. These endonucleases show high DNA degradation activity in blood, which is considered a normal physiological phenomenon whose purpose is to maintain homeostasis in the body and to protect against viruses [15,16]. However, the kinetics of long DNA fragmentation by serum nucleases and its role in the appearance of chromatosome-associated cirDNA fragments in the blood have yet to be fully elucidated.

NET were discovered in 2004 [17], and have been presented as a potential bactericidal killing mechanism. Their mechanism has been described as involving the activation of neutrophils to release their DNA, which is laden with powerful enzymes (such as neutrophil elastase and myeloperoxidase), that traps and kills microbes [18]. Interestingly, NET are found in the same pathological conditions where high concentrations of cirDNA have been reported, such as autoimmune diseases [19], inflammatory diseases [20], sepsis [21], thrombotic illnesses [22], and cancer [23–25]. However, despite the clinical potential inherent in such observations, no report exists which delineates the mechanistic process by which genomic DNA from NET is degraded into circulating mononucleosomes. Our studies now address that. Specifically, we believe that chromatin fragments, oligonucleosomes and nucleosomes can be liberated by the process of NET degradation which occurs during inflammation, and that these contribute to the pool of cirDNA and histones in blood [25]. We have already demonstrated the association of NET formation with cirDNA production in metastatic colorectal cancer [26], and we also postulate that NET have a key role in COVID-19 pathogenesis [25].

The fragmentation of cirDNA and NET-associated biomarkers is of particular interest in identifying the cirDNA’s cell/tissue-of-origin, and as such could have numerous clinical applications. It has also been shown that upon activation, neutrophil elastase (NE) escapes from azurophilic granules and translocates to the nucleus, where it partially degrades specific histones, promoting chromatin decondensation. Subsequently, myeloperoxidase (MPO) synergizes with NE in driving chromatin decondensation independent of its enzymatic activity [27]. These data led us to suggest that NET associated enzymes could modify the cirDNA structure and, thus change its fragment size distribution in patients with NET associated disorders. We have demonstrated the particular fragmentation of cirDNA in the plasma of cancer patients, and were the first to postulate that cirDNA fragment size profile (fragmentomics, [28]) could be used to distinguish the cirDNA of cancer and healthy subjects [29,30]. Our recent fragmentomics studies based on shallow whole genome sequencing (sWGS) revealed the detailed cirDNA size profiles associated with chromatosome structures in the plasma of healthy individuals (HI) and cancer patients [31]. Thus, cirDNA fragmentation pattern and NET associated biomarkers are being studied as a possible means of stratifying individuals [29–35].

In the present study (Fig. 1), we used the synergistic analytical information provided by qPCR, sWGS, and capillary electrophoresis to unequivocally observe the following: the kinetics of genomic DNA degradation by serum nucleases *in vitro*; the production of extracellular DNA (or cfDNA) by *ex vivo* activated neutrophils; the *in vitro* NET degradation by serum nucleases. This enabled us to precisely measure cirDNA size over a wide range of lengths, thus obtaining information about DNA structures which can be detected during the processes of NET production and degradation. We also performed an *in vivo* study in knockout mice, and an *in vitro* study of genomic high molecular weight (gHMW) DNA degradation, in order to elucidate the role of NE and MPO in cirDNA fragmentation. We then compared the NET associated markers and fragmentation size profiles of cirDNA in plasma obtained from patients with COVID-19, systemic lupus erythematosus (SLE) patients, metastatic colorectal cancer (mCRC) patients and healthy individuals (HI).

**Fig. 1:**
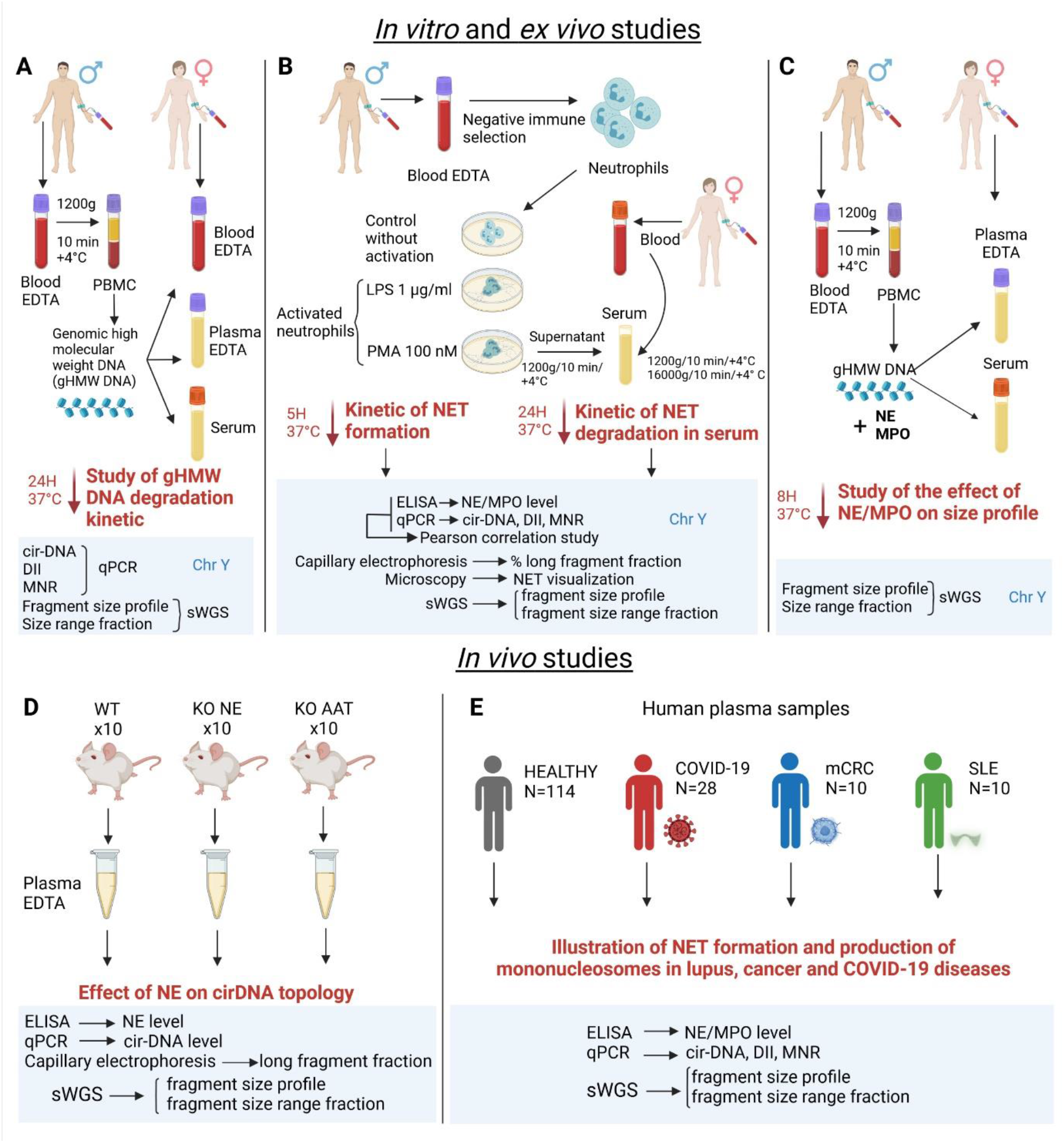
Schematic overview of the *in vitro, ex vivo* and *in vivo* studies of cirDNA production by NET. (A) *In vitro* study of gHMW DNA degradation in blood fluids using Y chromosome analysis. (B) *Ex vivo* study of NET production and degradation using Y chromosome analysis. (C) *In vitro* study of the effect of NE/MPO on cfDNA size profile using Y chromosome analysis. (D) *In vivo* study of the effect of NE on cirDNA topology. (E) *In vivo* study of NET associated disorders.

## Methods

### Patients and healthy individuals

Plasma samples from 28 patients with severe COVID-19 (at the entrance of Intensive Care Unit) were provided by the CHU hospital of Montpellier (Centre Hospitalier Universitaire de Montpellier, France); approval number assigned by the IRB: IRB-MTP_2021_01_202100715. We also included 10 mCRC patients from the screening procedure of the ongoing UCGI 28 PANIRINOX study (NCT02980510/EudraCT n°2016-001490-33). Plasma samples from 10 female patients with systemic lupus erythematosus (SLE) were provided by Dr. P. Simon from the Department of Sports Medicine, Prevention and Rehabilitation of Johannes Gutenberg University (Mainz, Germany). The study was approved by the Human Ethics Committee of Rhineland-Palatinate, Germany (number 2018-13039) and conformed to the standards of the Declaration of Helsinki of the World Medical Association and was registered under ClinicalTrials.gov Identifier: NCT03942718. We also analyzed 114 healthy individuals (HI, 59 men and 55 women) from the Etablissement Français du Sang (EFS), which is Montpellier’s blood transfusion center (Convention EFS-PM N° 21PLER2015-0013). These samples were initially screened (virology, serology, immunology, blood numeration) and ruled out wherever any abnormality was detected.

### Animals

Animal experiments involving WT, NE and AAT KO mice corresponded to the European guideline 2010/63/EU, and were authorized by the Lower Saxony State Office for Consumer Protection and Food Safety, Wardenburg, Germany (permission 18/2894).

### Extracellular DNA nomenclature

We recently published the first article proposing a standard nomenclature to be used in the cirDNA field, to homogenize the various terminologies currently in use [1]. Thus, we clearly differentiate cell-free DNA (cfDNA), which can be found in various biological sources (cell culture supernatant, blood, urine, pap smears, LCR …), from circulating DNA, which is found only in blood or lymph nodes. While there both groups share numerous characteristics, their physiological localization results in clear differences, as shown for instance by their fragment size profile (i.e. blood vs urine) [1]. In this study, we clearly differentiate cfDNA, detected in the gHMW DNA or NET *in vitro* degradation experiments, from cirDNA, detected in plasma from human or mice subjects.

### gHMW DNA preparation

A blood sample from a male healthy individual was collected in a 10 ml EDTA tube. After centrifugation of the blood at 1200 g for 10 min at +4°C, the PBMC layer was collected into a separate tube. The PBMCs were washed twice with PBS and pelleted by centrifugation at 800 g. The supernatant was discarded, and the cell pellet resuspended in 1 ml of PBS with the addition of a protease inhibitor cocktail (cOmplete™ Protease Inhibitor Cocktail, Roche) in order to protect histones from degradation by proteases during the next step of cell lysis. The cells were lysed by 10 freeze-thaw cycles. The lysate was centrifuged at 800 g for 10 min at +4°C, in order to precipitate the non-lysed cells. The supernatant of this cell extract containing a high molecular weight genomic DNA (gHMW DNA, chromatin) was collected and used for the following experiments on degradation of DNA in serum and plasma. The 25 µL aliquot of the cell extract was used for quality and quantity control of chromatin in the cell extract. The DNA was extracted using a QIAamp DNA blood mini kit (Qiagen). Its quantity and integrity were analyzed by: (1) qPCR, using the two pairs of primers for amplification of the short and long fragments (67 bp and 320 target on *KRAS* gene); and by (2) capillary electrophoresis on Fragment Analyzer (Agilent, HS Large Fragment Kit DNF-493). The quality control of gHMW DNA before degradation showed a DNA concentration of 10 ng/µL, and DNA fragments ranging from 2 - 40 kb, peaking at 12 kb. (Additional file 1: Fig. S1).

### Plasma/serum isolation

Samples were handled according to pre-analytical guidelines previously established by our group [36]. Blood samples from HI or patients were collected in 10 ml EDTA tubes (K2E, REF 367525, BD Vacutainer) for plasma or in 4 ml CAT tubes (Clot Activator Tube, REF 369032, BD Vacutainer) for serum. Serum was used only for the experiments involving gHMW DNA or NET degradation. Blood from mCRC patients (n=10) was collected in STRECK tubes (Cell-Free DNA BCT®) and was sent within 24H of collection at room temperature from the recruiting institutions to our laboratory (IRCM, Institut de Recherche en Cancérologie de Montpellier, U1194 INSERM). The blood was then centrifuged at 1200 g at 4°C for 10 min. The supernatants were isolated in sterile 1.5 mL Eppendorf tubes and centrifuged at 16000 g at 4°C for 10 min [29]. Afterwards, the plasma or serum was either immediately used for the experiment on gHMW DNA or NET degradation, or was stored at -20°C until cirDNA extraction or ELISA analysis.

### Design of experiment for the study of gHMW DNA and NET degradation

The study of gHMW DNA and NET degradation was done by incubation of the chromatin from PBMC lysate in blood fluids. We used gHMW DNA from a male donor, and blood/plasma/serum from a female donor, to further analyze Y chromosome fragments that belong exclusively to the gHMW DNA under study, and which obviously are absent in the plasma of the female donor. The size profile of the degraded Y chromosome was studied by qPCR and shallow WGS.

### Degradation of gHMW DNA in blood fluids

The 20 µl of chromatin equivalent to 200 ng of genomic DNA was added to 5 aliquots of 500 µL of blood EDTA, plasma EDTA or serum. These samples were incubated at 37 °C for 0 min, 30 min, 2H, 8H and 24H. After incubation, the plasma was isolated from blood EDTA as described above. Then, cfDNA was extracted from 0.5 ml of plasma or serum using the QIAmp DNA Mini Blood kit (Qiagen, CA), according to the ‘‘Blood and body fluid protocol’’ and our own detailed protocol [37]. CfDNA samples were kept at 20 °C until use. Due to EDTA associated calcium chelation inhibitory effect on nucleases, comparison of plasma vs serum incubation may help in estimating nuclease role in degrading gHMW DNA.

### Degradation of gHMW DNA in blood fluids in presence of NE and MPO

The 50 µl of chromatin equivalent to 500 ng of genomic DNA was added to 1mL of plasma EDTA or serum. 200 ng of NE (Sigma, 324681) or 200 ng MPO (Sigma, 475911) or both was added to the samples. Control samples were incubated under the same conditions, but without the addition of NE or MPO. These samples were incubated at 37 °C for 2H and 8H. Then, cfDNA was extracted from plasma or serum using the QIAmp DNA Mini Blood kit (Qiagen, CA), according to the ‘‘Blood and body fluid protocol’’ and our own detailed protocol [37]. CfDNA samples were kept at 20 °C until use.

By subtracting values obtained in medium with NE/MPO from those obtained in the controls, we are able to estimate the DNA degradation rate in various conditions. Based on the strong inhibitory effect of EDTA on nuclease activity (95% and ∼85% of DNA remain following 2 and 8 hours, respectively), the overall, independent nuclease activity in serum is estimated by subtracting the DNA concentration in control serum from that found in control plasma over time. Assuming that overall DNA degradation in serum is due to the combined activities of nuclease and of NE/MPO, we estimated the degradation rate deriving specifically from NE, MPO, and NE plus MPO by subtracting the DNA concentration in serum to which NE and/or MPO has been added from that found in control serum over time. Overall degradation rates are expressed as ng DNA/mL/minutes, and cannot be compared to an enzymatic activity.

### NET induction

For neutrophil preparation, blood from a healthy male volunteer was collected into two 10 ml EDTA tubes using standard phlebotomy techniques. Neutrophils were isolated from 20 ml of blood with EasySep™ Human Neutrophil Isolation Kit (STEMCELL Technologies, British Columbia, Canada, 19666) according to the manufacturer’s instructions. The neutrophils were quantified using Trypan Blue staining on Countess II automated cell counter. Cell suspension was diluted to the cell concentration 5.6 × 10^5^ cells/ml with the RPMI medium containing 10% of serum from the donor of neutrophils. The neutrophils were seeded (5.6 × 10^5^ cells/ml; 1.1 million cells per well) in the 6-well plates (Falcon, REF 353046). The neutrophils were incubated at 37°C for 30 min, 1H, 2H, 4H and 5H under the following conditions: control (non-stimulated neutrophils), neutrophils stimulated by PMA (PHORBOL 12-MYRISTATE 13-ACETATE, Sigma, P8139-1MG) in concentration 100 nM, and neutrophils stimulated by LPS (LIPOPOLYSACCHARIDES FROM ESCHERICHIA COLI, Sigma, L4391-1MG) in concentration 1 µg/ml. All experiments were performed in duplicate. After incubation, cell media was aspirated and centrifuged at 1200 g for 10 min at +4 °C in order to collect cell-free media.

### Degradation of NET in blood fluids

As part of our study, we also investigated the kinetics of NET degradation by serum nucleases *in vitro* by incubation of the supernatant of PMA stimulated (5H) male neutrophils from the previous part of the study in a fresh female serum. We also used the X/Y differentiation to distinguish NET-derived male DNA from female cirDNA derived from serum. The qPCR analysis was performed targeting the *SRY* gene on the Y chromosome, and sWGS results took into consideration only reads aligning on the Y chromosome.

The 200 µl of supernatant of the PMA stimulated neutrophils (5H) equivalent to 50 ng of genomic DNA was added to aliquots of 500 µL of plasma or serum. These samples were incubated at 37 °C for 5 min, 10 min, 20 min, 30 min, 2H, 8H, and 24H for NET degradation in serum; with 24H incubation in plasma used as a control. After incubation, cfDNA was extracted from 0.5 ml of plasma/serum using the QIAmp DNA Mini Blood kit (Qiagen, CA), performed according to the ‘‘Blood and body fluid protocol’’ and our own detailed protocol [37]. CfDNA samples were kept at -20 °C until use.

### DNA extraction from plasma

CirDNA from plasma, serum or cell medium was extracted using the QIAmp DNA Mini Blood kit (Qiagen), used according to the ‘‘Blood and body fluid protocol’’, using 80 µl of elution volume. DNA extracts were kept at -20 °C until used.

### Quantification of DNA by qPCR

The concentration of cf- or cirDNA was assessed using an integrated PCR system that targeted sequences within the same region. The designed amplification sizes were 73 bp and 246 bp targeted *SRY* gene on the Y chromosome for cfDNA quantification; and 67 bp and 320 bp targeted *KRAS* gene for cirDNA quantification. This primer construction enabled the calculation of a DNA integrity index (DII) by calculating the ratio between the concentrations obtained with the primer pairs, by targeting both a long (246 bp or 320 bp) and a short (73 bp or 67 bp) sequence. Targeting long sequences (over 250 bp) are unable to amplify cirDNA fragments packaged in mononucleosome, while targeting short sequences (60-80bp) amplifies almost all DNA fragments, including those associated with mononucleosomes (99% of cirDNA fragments from HI are detected by targeting a 60bp sequence, [31]) qPCR amplifications were carried out at least in duplicate in a 25 µL reaction volume on a CFX96 instrument using the CFX manager software (Bio-Rad). Each PCR reaction mixture was composed of 12.5 µl PCR mix (Bio-Rad Sso advanced mix SYBR Green), 2.5 µl of each amplification primer (0.3pmol/ml, final concentration), 2.5 µl PCR-analyzed water, and 5 µl DNA extract. Thermal cycling consisted of three repeated steps: a 3 min Hot-start Polymerase activation and denaturation step at 95 °C, followed by 40 repeated cycles at 95 °C for 10 s, and then at 60 °C for 30 s. Melting curves were obtained by increasing the temperature from 55 °C to 90 °C, with a plate reading every 0.2 °C. The concentration was calculated from Cq detected by qPCR, and also a control standard curve on genomic DNA of known concentration and copy number. Serial dilutions of genomic DNA from human placenta (G1471, Promega) were used as a standard for quantification and their concentration.

### DNA analysis by capillary electrophoresis

The analysis of DNA by capillary electrophoresis was done using the Fragment Analyzer 5200 (Agilent) and DNF-493 kit, according to manufacturer’s instructions.

### Library preparation for shallow WGS

In this study, we analyzed DNA size profiles using shallow WGS, which allows the accurate estimation of the number of fragments of a certain length with a resolution of 1 bp. DSP libraries were prepared with the NEBNext® Ultra™ II kit. For library preparation, a minimum of 1 ng of cf- or cirDNA was engaged without fragmentation, and recommendations from kit providers were followed. Briefly, for DSP with the NEB kit, Illumina paired-end adaptor oligonucleotides were ligated on repaired A-tailed fragments, then SPRI purified and enriched by 11 PCR cycles with UDI primers indexing, and then SPRI purified again. The SPRI purification was adjusted to keep the small fragments around 70b of insert. Finally, the libraries to be sequenced were then precisely quantified by qPCR, to ensure that the appropriate quantity was loaded on to the Illumina sequencer, in order to obtain a minimum of 1.5 million of clusters.

### Analysis of cirDNA by shallow WGS

All libraries were sequenced on MiSeq 500 or NovaSeq (Illumina) as Paired-end 100 reads. Image analysis and base calling was performed using Illumina Real Time Analysis with default parameters. The individual barcoded paired-end reads (PE) were trimmed with Cutadapt v1.10 to remove the adapters and discard trimmed reads shorter than 20 bp Trimmed fastq Files were aligned to the human reference genome (GRCH38) using the MEM algorithm in the Burrows-Wheeler Aligner (BWA) v0.7.15. The insert sizes were then extracted from the aligned bam files with the TLEN column for all pairs of reads having an insert size between 0 - 1000 bp. Supplementary alignments, PCR duplicates and reads with MapQ<20 were filtered out from the analysis. Summary statistics of sWGS data is presented in Additional file 1 (Table 1S).

### Human Myeloperoxidase (MPO) and Neutrophil Elastase (NE) quantifications

MPO and NE concentrations were measured using enzyme-linked immunosorbent assay (ELISA), according to the manufacturer’s standard protocol (Duoset R&D Systems, DY008, DY3174, and DY9167-05). Briefly, captured antibodies were diluted at the working concentrations in the Reagent Diluent (RD) provided in ancillary reagent kits (DY008), and coated overnight at room temperature (RT) on 96-well microplates with 100 μL per well. Then, captured antibodies were removed from the microplates, and wells were washed three times with 300 μL of Wash Buffer (WB). Microplates were blocked at RT for 2H by adding 300 μL of RD to each well. RD were removed from the microplates, and wells were washed three times with 300 μL of WB. Then, 100 μL of negative controls, standards and plasma samples (diluted 1/10) were added to the appropriate wells for 1H at RT. Samples, controls and standards were removed from the microplates, and wells were washed three times with 300 μL of WB. Detection antibodies were diluted at the working concentrations in the RD, then added by 100 μL per well, for 1H at RT. Detection antibodies were removed from the microplates, and wells were washed three times with 300 μL of WB. Then, 100 μL of Streptavidin-HRP was added to each well and microplates were incubated at RT for 30 min. The wash was repeated three times. Finally, 100 μL per well of substrate solution was added and incubated for 15 min, and the Optical Density (OD) of each well was read immediately at 450 nm with the PHERAstar FS instrument using the PHERAstar control software.

### Statistical Analysis

Statistical analysis was performed using the GraphPad Prism V6.01 software. Correlation analysis was performed using the Pearson test. The Mann-Whitney test was used to compare means. A probability of less than 0.05 was considered to be statistically significant; *p<0.05, **p<0.01; ***p< 0.001; ****p< 0.0001.

## Results

### Kinetics of gHMW DNA degradation in blood fluids

To study the degradation of gHMW DNA, we examined nuclear chromatin from PBMC during incubation in blood fluids. We observed a progressive decrease of total DNA concentration and long DNA fragments (>246 bp) over the course of 24 hours’ incubation in serum (Fig. 2A), whereas no change in terms of concentration and DII occurred to DNA from full blood or plasma collected in EDTA tubes (Fig. 2A). This showed that shorter fragments shorter than 246 bp did not exist or existed only in minimal quantities before incubation in serum, in contrast to during incubation in serum. More details of the results of this experiment are provided in Additional file 1 (Fig. S5)

**Fig. 2:**
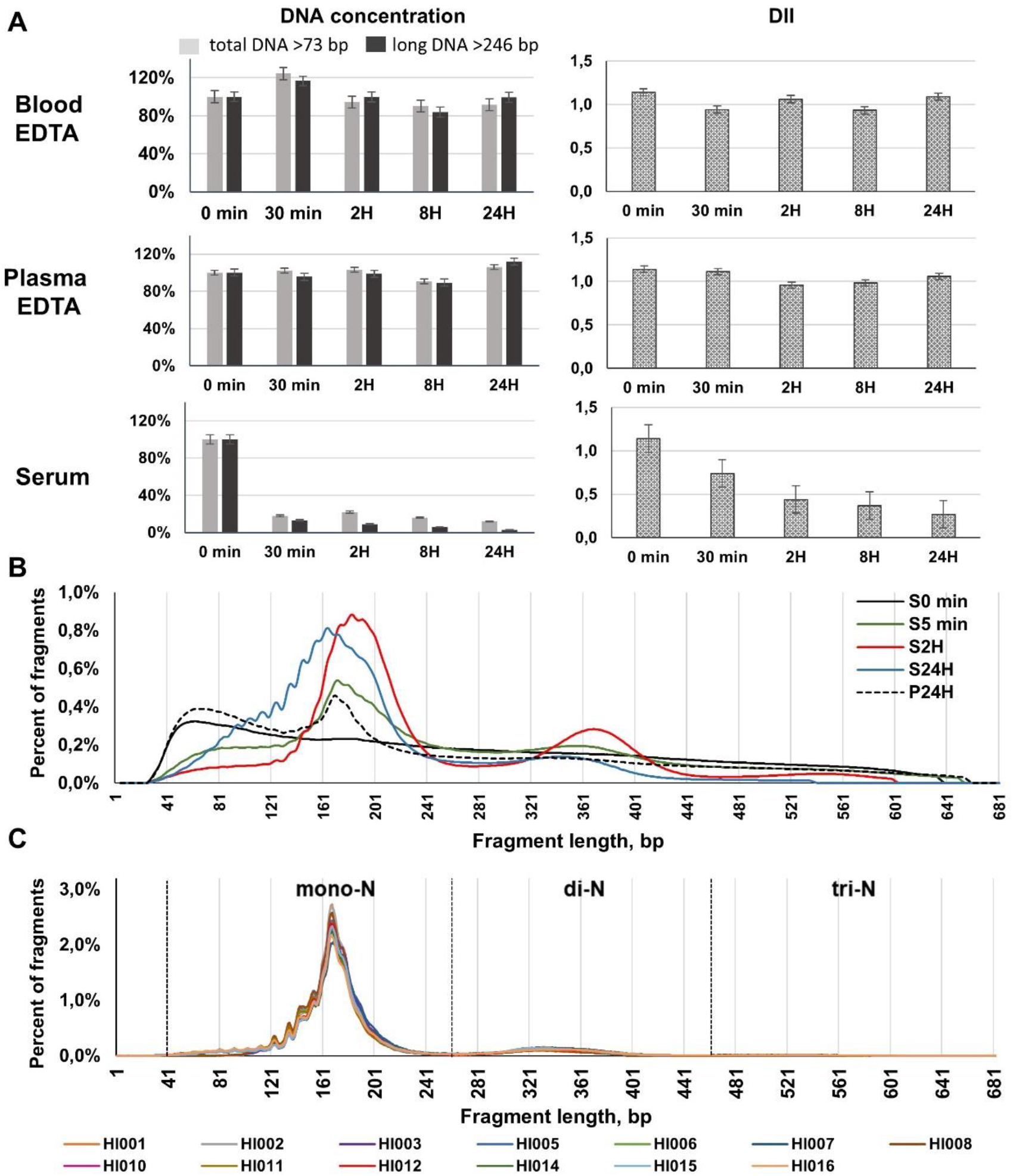
Kinetics of gHMW DNA degradation in blood fluids. (A) The kinetics of total and long DNA concentrations, as determined by qPCR, presented as a fraction of the initial concentration and DII (DNA integrity index) following gHMW DNA incubation up to 24 hours at 37°C in blood EDTA, plasma EDTA and serum. Data are represented as mean ± SEM. (B) Size profiles of gHMW DNA samples before incubation in blood fluids (S0) and after incubation at 37°C in plasma for 24 hours (P24H) or in serum for 5 minutes (S5 min), 2 hours (S2H) or 24 hours (S24H) as determined by sWGS. (C) CirDNA fragment size profile of the thirteen healthy subjects, as determined by sWGS. Examination of gHMW DNA degradation and size analysis were performed and size profile using Y chromosome analysis.

While no peak appears at the start of incubation, sWGS revealed as soon as following 5 minutes incubation, the appearance of the peaks at 167 bp and 350 bp (Fig.2B) which correspond to mono-N DNA and di-N DNA as found in cirDNA size profile of healthy individuals (Fig. 2C). The DNA size profile after 24 hours’ incubation in serum showed an accumulation of short fragments below 160 bp (down to 41 bp), with the appearance of sub peaks with a ∼10 bp periodicity revealing at that size range mono-nucleosome footprints (Fig.2B). Thus, mono-N DNA accumulated up to 81.3% (Additional file 1, Table 2S) during the 8H incubation period in the 20-1000 bp size range. Whereas mono-, di- and tri-N pattern appeared following 5 minutes incubation, di-N and tri-N DNA fractions decreased from 2 to 24 hours incubation (Fig 2B, Additional file 1,Table 2S)

The use of serum shows here several limitations, given that the preparation of serum may lead to some cell degradation and possibly to the extracellular release of DNase or DNA, and the apparition of components linked to coagulation. While extracellularly released intracellular DNase should be diluted in a large amount of serum nuclease, and extracellular DNA release cannot interfere due to the specific quantification of Chr Y sequences, a bias may emerge as we cannot exclude the possibility of coagulation derived components interfering with the DNA degradation rate, by interfering with DNA stability, for instance. Thus, the serum in which DNA degradation is examined is artificial, limiting the direct comparison of these observations to what occurs to the plasma in blood circulation.

### *Kinetics of* ex vivo *NET production*

*Ex vivo* isolated neutrophils were stimulated by PMA or LPS, in order to investigate NET associated DNA release (Fig. 3). The qPCR analysis revealed a sharp increase of cf-nDNA and cf-mtDNA concentrations in the supernatant of stimulated neutrophils, with minor increase in control cells (Fig. 3A,B). After 5 hours’ incubation, the total cfDNA concentrations were 7 - 35 fold higher in the supernatant of LPS- and PMA-stimulated neutrophils than in the supernatant of non-stimulated controls. NE and MPO concentration, increased up to 5-fold in the supernatant of stimulated neutrophils as compared to unstimulated control after 5 hours incubation in cell culture (Fig. 3D). The Pearson correlation study revealed strong positive correlations between DNA and NET protein (NE, MPO) markers in stimulated neutrophils (Additional file 1, Fig. S4). The kinetics of the release in the cell culture supernatant of MPO, NE, cf-nDNA and cf-mtDNA are all remarkably similar up to 5H incubation, suggesting the co-occuring release of these components, and consequently suggesting that cfDNA may be a byproduct of NET.

**Fig. 3:**
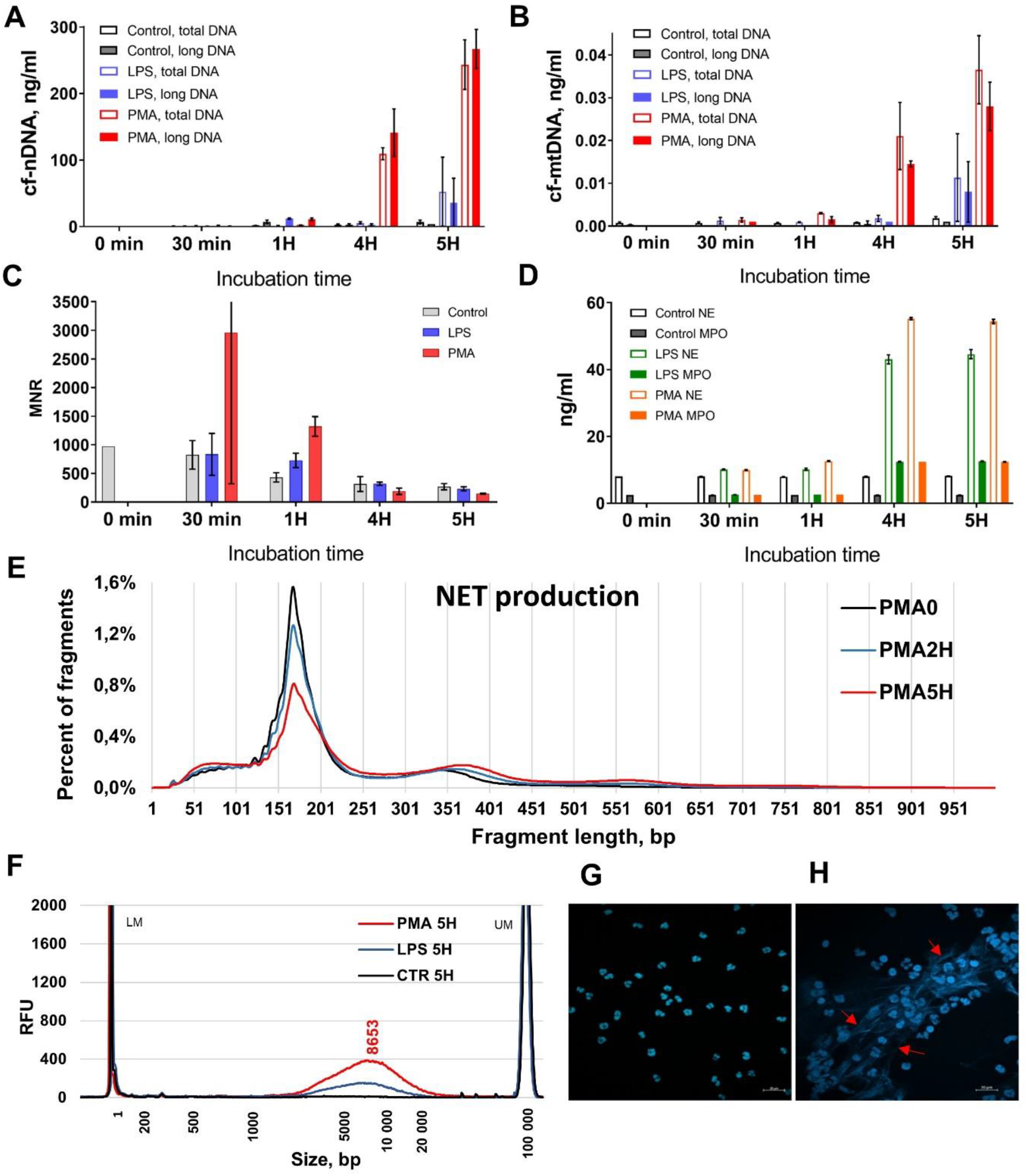
Kinetics of NET production *in vitro*. The kinetics of cfDNA, as determined by qPCR, in the supernatant of control non-stimulated and stimulated by 1 mg/ml LPS or 100 nM PMA human neutrophils from a healthy donor following incubation for up to 5 hours at 37°C in cell culture: (A) total and long (>320 bp) nuclear cf-nDNA concentration; (B) total and long (>310 bp) mitochondrial cf-mtDNA concentration; (C) MNR (ratio of mitochondrial to nuclear DNA concentration). (D) Kinetics of NE and MPO in the supernatant of the same control non-stimulated and stimulated by 1 mg/ml LPS or 100 nM PMA. Data are represented as mean ± SD. (E) NET production: the cfDNA size profiles of the supernatant of human neutrophils before incubation and stimulation (PMA 0); and following stimulation by PMA 100 nM and incubation at 37°C in cell culture for 2 hours (PMA2H) and 5 hours (PMA5H), as determined by sWGS. (F) Analysis of cfDNA size profile by capillary electrophoresis (Fragment Analyzer) in the supernatant of control non-stimulated (CTR 5H) and stimulated by LPS (LPS 5H) and PMA (PMA 5H) neutrophils following 5 hours’ incubation in cell culture. (G,H) The fluorescent microscopic image of Hoechst stained neutrophils stimulated by LPS (H) and control neutrophils (G) following 5 hours’ incubation in cell culture. Examination of DNA content using Y chromosome analysis.

Fragment size profile analysis by sWGS of DNA from the supernatant of *ex vivo* stimulated neutrophils (Fig. 3E) revealed the typical chromatin organization pattern as observed for cirDNA derived from human blood (Additional file: Fig. S3). Overall, there was a progressive increase of di-N, tri-N and long DNA associated peaks (Fig. 3E; PMA 2H, PMA 5H), as compared to baseline (PMA0) (Fig. 3E). Notably, while the mono-N population peaked at 167 bp irrespective of incubation time, the di-N DNA peaked at 365 bp and at 341 bp in PMA 5H and baseline (PMA0), respectively. In addition, PMA 5H stimulated neutrophils supernatant showed a small third peak at 560 bp, corresponding to tri-N DNA. Capillary electrophoretic analysis demonstrated that activated neutrophils produced HMW DNA whose mean size was 8,600 bp. In contrast, non-stimulated neutrophils did not show an accumulation of long DNA fragments even after 5H of incubation in cell culture (Fig. 3F).

To visualize the NET structure, we performed a DNA specific Hoechst staining of stimulated (Fig. 3H) and control neutrophils, (Fig. 3G) with presence of NET fibers observed around the stimulated neutrophils.

### Kinetics of in vitro NET degradation in blood

To study NET degradation, we placed the culture medium of 5H PMA-stimulated neutrophil obtained from a male donor in the plasma and serum of a healthy female donor. The concentrations of both total cfDNA and long cfDNA fragments decreased progressively in serum, with the cf-nDNA concentration after 24H incubation in serum being 3 - 4 fold lower than the cfDNA concentration at baseline (S0) (Fig. 4A). NET were seen to undergo rapid and strong degradation, with a significant loss of 28%, 48%, and 71% of NET derived DNA input being observed after 10 min, 30 min and 24H. The DII decreased from 1.0 to 0.6 during the first 30 min, then stabilized up to 24H (Additional file 1: Fig. S5). Incubation of NET in plasma for 24H did not lead to a significant change of the concentrations of total DNA or long DNA fragments (Fig. 4A), or of the DII (Additional file 1: Fig. S5). Capillary electrophoresis demonstrated the shortening and degradation of long DNA fragments during 8h incubation in serum: the average size of long DNA fragments decreased from 8782 bp to 2589 bp, with the fraction of DNA fragments in the 1K-10K bp range decreasing from 86% to 36% (Fig. 4B). Note, only <2% and 1% of this fragment length population remained following 2 and 24H incubation.

**Fig. 4:**
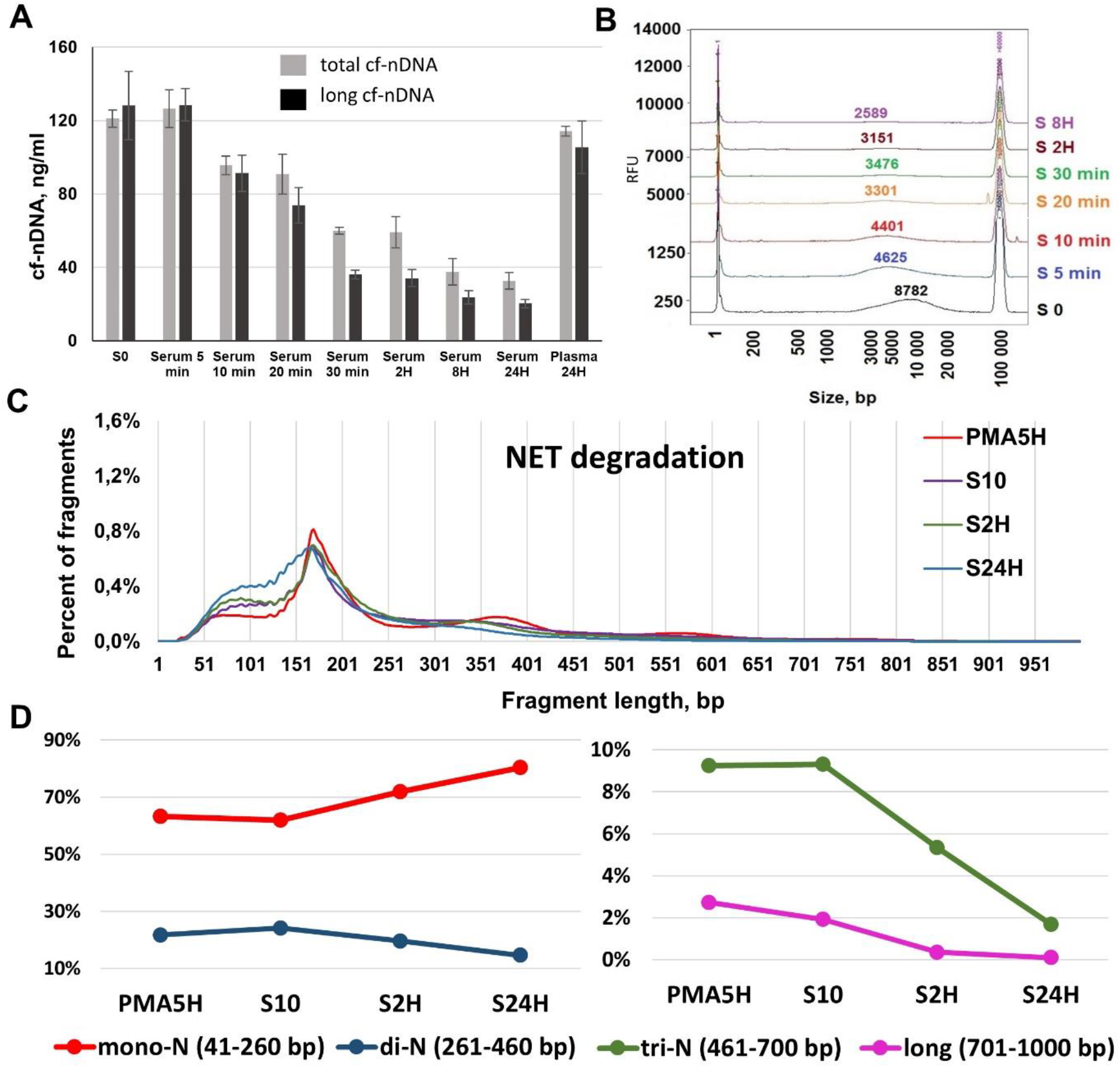
Kinetics of NET degradation in blood fluids. Kinetics of total and long (>246 bp) nuclear cell-free DNA (cf-nDNA) concentration following the degradation of NET (supernatant of PMA activated neutrophils) by incubation in serum/plasma up to 24 hours: before incubation (S0) and after incubation at 37°C in plasma for 24 hours or in serum for 5 min/10 min/20 min/30 min/2 hours/8 hours/24 hours. Data are represented as mean ± SD. (B) Analysis of cfDNA size profile by capillary electrophoresis (Fragment Analyzer) in the supernatant of PMA stimulated (5 hours) neutrophils (S0) and following its incubation in serum (S 5 min/S 10 min/S 20 min/S 30 min/S2H/S8H). (C) NET degradation in serum: the cfDNA size profiles of the supernatant of PMA stimulated (5 hours) neutrophils (PMA5H); and following incubation of this supernatant in serum at 37°C for 10 minutes (S10), 2 hours (S2H) and 24 hours (S24H). (E) Analysis of sWGS data for assessment of the cfDNA fractions corresponding to mono-, di- and tri-nucleosome associated fragments during NET degradation. Examination of DNA degradation and of size profile were performed using Y chromosome

sWGS fragment size profile analysis of NET-derived DNA, performed in the course of incubation in serum, revealed an evolution of the organization pattern of cfDNA associated chromatin. We observed a progressive accumulation of DNA fragments <167 bp, and a decrease of the di-N associated peak (260 - 460 bp) from 10 min to 24H incubation in serum (Fig. 4C). In contrast to the effect on NET production in culture medium (Fig. 3E), NET degradation in serum produced a shortening of the di-N DNA, with the maximum associated peak decreasing from 365 to 321 bp (Fig. 4C). Furthermore, the tri-N DNA fragment population disappeared following 24H incubation in serum (Fig. 4C).

When comparing the size profile analysis of DNA extracted during the incubation of stimulated neutrophils (NET production) and of DNA extracted in the course of incubation in blood fluids (NET degradation), we observed that (i), NET production was associated with the increase of fractions of mono-N, di-N, tri-N and long DNA fragments up to 65%, 22%, 9% and 2.7%, respectively. In contrast, NET degradation in serum led to an accumulation of the mono-N DNA fraction representing 80%, and a decrease of the di-N, tri-N and long DNA fractions down to 15%, 1.7% and 2.73% to 0.10%, respectively (Fig. 4D). The high, intermediate and low degradation rates of long, tri-N and di-N DNA is in contrast to the increase of mono-N DNA, suggesting that DNA in oligonucleosomes are degraded to mono-N DNA. The mono-N DNA amount corresponds to a balance between the respective rates of their degradation and their accumulation due to DNA lower nuclease sensitization when packed in mononucleosomes. When combining capillary electrophoresis and sWGS data, it appears that high molecular weight DNA from NET chromatin fibers are highly degraded, resulting in the indirect detection of oligonucleosomes, principally mononucleosomes.

### gHMW DNA degradation in presence of NE and MPO

As quantified by qPCR, after 2H incubation in serum containing NE or MPO, the amount of gHMW DNA showed a 3- and 5-fold decrease, respectively, as compared with control incubation (64 ng/ml and 35 ng/ml vs 207 ng/ml, respectively) (Fig. 5A).

**Fig. 5:**
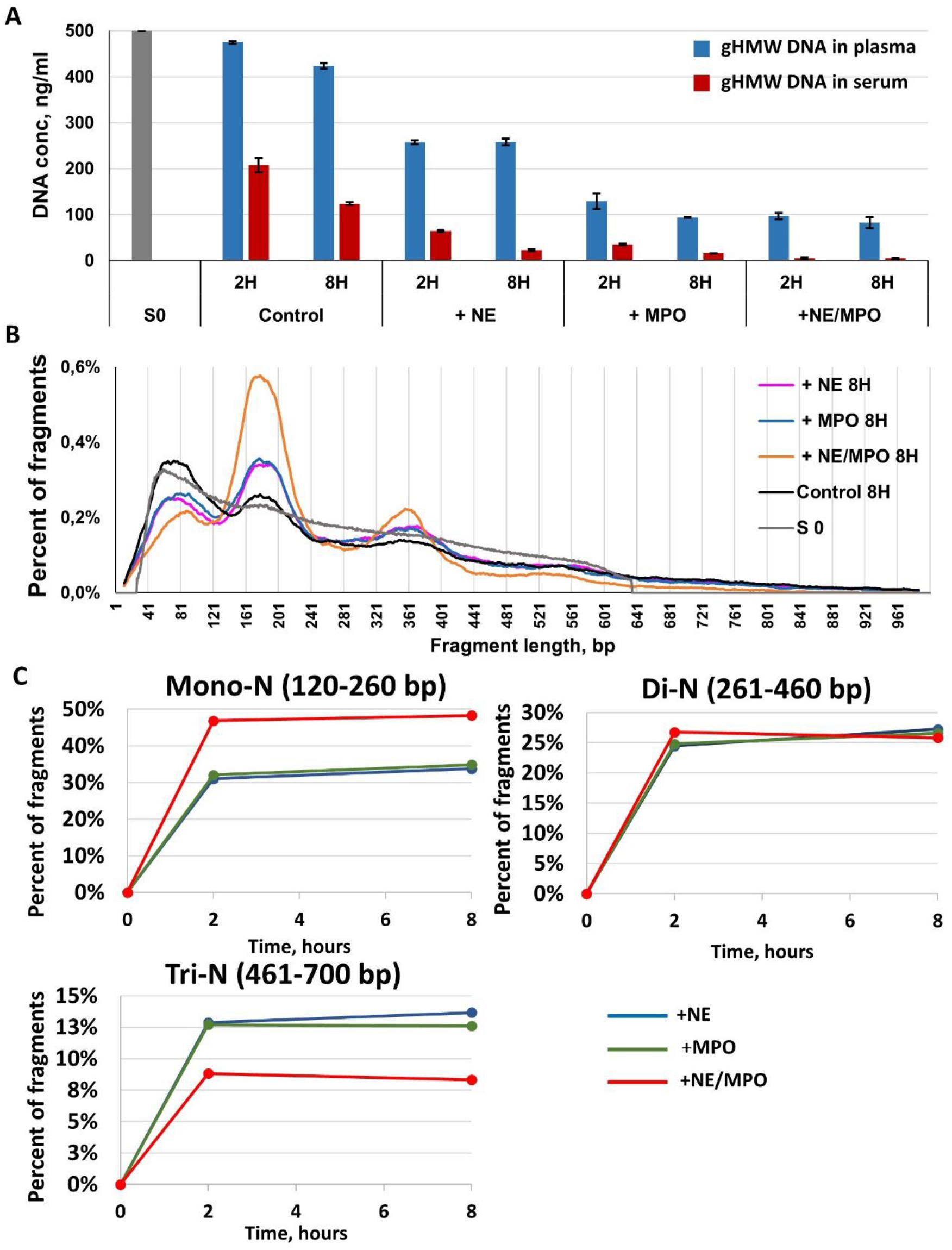
gHMW DNA degradation in presence of NE and MPO. (A) Quantification of total DNA by qPCR in cell lysate containing gHMW DNA before degradation (S0) and following gHMW DNA incubation at 37°C for 2 or 8 hours: in plasma or serum (control) and in plasma or serum in the presence of neutrophil elastase (+NE), myeloperoxidase (+MPO) or both (+NE/MPO). (B) CfDNA size profiles for gHMW DNA samples before incubation in blood fluids (S0) and following 8 hours of incubation at 37°C in serum: control 8 hours or in the presence of neutrophil elastase (+NE), myeloperoxidase (+MPO) or both (+NE/MPO). (C) Analysis of cfDNA fractions corresponding to mono-, di- and tri-nucleosome (Mono-N, Di-N, and Tri-N, respectively) associated fragments as determined by sWGS: cfDNA fractions in the course of gHMW DNA degradation in the presence of neutrophil elastase (+NE), myeloperoxidase (+MPO) or both (+NE/MPO). Examination of gHMW DNA degradation and of size profile were performed using Y chromosome analysis.

We observed an even more significant difference after 8h incubation, as the presence of NE and MPO led to 5.4-fold and 8-fold decrease, respectively, of total DNA concentration, as compared to control incubation (23 ng/ml and 16 ng/ml vs 124 ng/ml) (Fig. 5A). These variations appeared even more impressive with the simultaneous addition of both NE and MPO, with 8h incubation leading to a 25-fold decrease of total DNA concentration, as compared to control incubation (5 ng/ml vs 124 ng/ml) (Fig. 5A). ∼99% of gHMW DNA were degraded following the later condition.

As shown in Fig. 5A and in Additional file 2 the estimated degradation rate within the first two hours and from 2 - 8 hours’ incubation is 11.6- and 4.9-fold higher in serum than in plasma, respectively. It appears that within the first two hours of incubation the degradation rate is 3- and 4-fold higher than that observed during the 2-8 hours incubation period for controls and NE/MPO containing serum. The estimated degradation rate in serum or in plasma is always significantly higher in NE/MPO containing serum than in controls, irrespective of whether the first 2H or 2-8H incubation periods are considered. For instance, data showed 1.5-, 1.6-, and 1.8-fold increase in NE, MPO and NE + MPO in serum, respectively; similarly, albeit to a larger extent, ∼10-, 15- and 16-fold increases were shown in NE, MPO and NE + MPO in plasma, respectively. Whereas the degradation rate is 11.6-fold higher in serum as compared to plasma in controls during the first two hours, it is much more similar when adding NE, MPO, and NE + MPO (1.8-, 1.3- and 1.2-fold, respectively). The highest estimated DNA degradation rate (4.12 ng/mL/minutes) was found in NE + MPO containing serum during the first two hours of incubation. Note, there is only a slight increase in the degradation rate in serum vs plasma during both incubation time periods when MPO and NE + MPO are added (∼1.2-fold in both 0-2H and 2-8H time periods, respectively), and when NE is supplemented to serum and plasma (∼1.8- and 2-fold, in 0-2H and 2-8H time periods, respectively). This is a striking demonstration of the nearly equivalent DNA degradation rate deriving from NE, and to a greater extent deriving from MPO activities, irrespective of nuclease inhibition. gHMW DNA degradation rate for each condition of incubation are described in Additional file 2.

From the data shown in Fig 5A, we can estimate the capacity of NE and/or MPO to effect the gHMW DNA degradation rate. Data suggest that: (i), the addition of NE and/or MPO to plasma or to serum greatly increases HMW DNA degradation (Fig. 5A); (ii), NE and MPO independently increase HMW DNA degradation; (iii), NE and/or MPO greatly increase DNA degradation in the presence of high nuclease activity as well as when nuclease activity is strongly inhibited; (iv) MPO shows an increase in DNA degradation approximately twice that of NE; and (v), NE and MPO may act separately to contribute to DNA degradation, while there is only a slight increase when NE and MPO are combined. The gHMW DNA degradation rate for each condition of incubation is described in Additional file 2.

We also observed a decrease of gHMW DNA during incubation in plasma. But these variation levels were not as pronounced as in the case of gHMW DNA incubation in serum (Fig. 5A).

The fragment frequency as determined from the size profile of gHMW DNA before incubation in serum decreased linearly from 50 to 700 bp (Fig. 5B), which was similar to what had been observed previously (Fig. 2B). After incubation in serum, the gHMW DNA fragment size profile showed the specific chromatin organization pattern with peaks corresponding to mono-N, di-N and (less prominently) tri-N peaks. All three peaks were more pronounced when gHMW DNA was degraded in the presence of NE or MPO than in serum only (control sample). Degradation with the combined incubation of both NE and MPO was associated with the most prominent peaks of mono-N and di-N (Fig. 5B). gHMW DNA incubation in serum for 8H (Control 8H) led to a clear modification of the size profile, as determined by sWGS. Incubation in serum with NE or MPO alone for either 2H or 8H provoked an equivalent increase of mono-N and di-N DNA populations. Incubation in serum with NE and MPO combined further enhanced the prominence of both corresponding peaks (Fig. 5B). As shown in Fig. 5C, the increase of the mono-N DNA fraction is faster within the first two hours of incubation than that observed during the 2-8 hours incubation period (Fig.5C), mirroring the higher cirDNA degradation rate estimated during the first two hours of incubation.

### In vivo study on the role of NE in cirDNA fragmentome

In order to elucidate the role of NE in DNA degradation, we performed an *in vivo* study of cirDNA in WT, NE KO, and alpha antitrypsin (AAT) KO mice [38]. EDTA blood samples were collected from these 3 groups of mice (10 mice in each group), and plasma was isolated for further NE and cirDNA analysis. Quantification of NE by ELISA confirmed the absence of NE in NE KO mice, and showed that the concentration of NE was significantly higher in AAT KO mice than in WT mice (2.52±0.39 ng/ml in WT vs 3.31±0.67 in AAT KO, p=0,005).

The cirDNA size profiles in plasma of WT, NE KO and AAT KO mice was determined by sWGS (Fig. 6A). As observed in human blood, the cirDNA fragment size profile of each group of mice (N=10) revealed the mammalian chromatin organization, with the main mono-N population peaking at 167 bp, and the second, less prominent population peaking at 335 - 355 bp, which corresponds to di-N. The WT and AAT KO mice plasma showed similar profiles, with a major population peaking at 167 bp and at subpeaks with ∼10 bp periodicity below 180 bp, and with a minor population peaking at 335 bp. The cirDNA size profile of NE KO mice differed from that of the WT and AAT mice, exhibiting a shift to a longer average length of cirDNA fragments. Detailed descriptions of the size profile characteristics of the three types of mice are presented in Additional file 1 (Fig. S2).

**Fig. 6:**
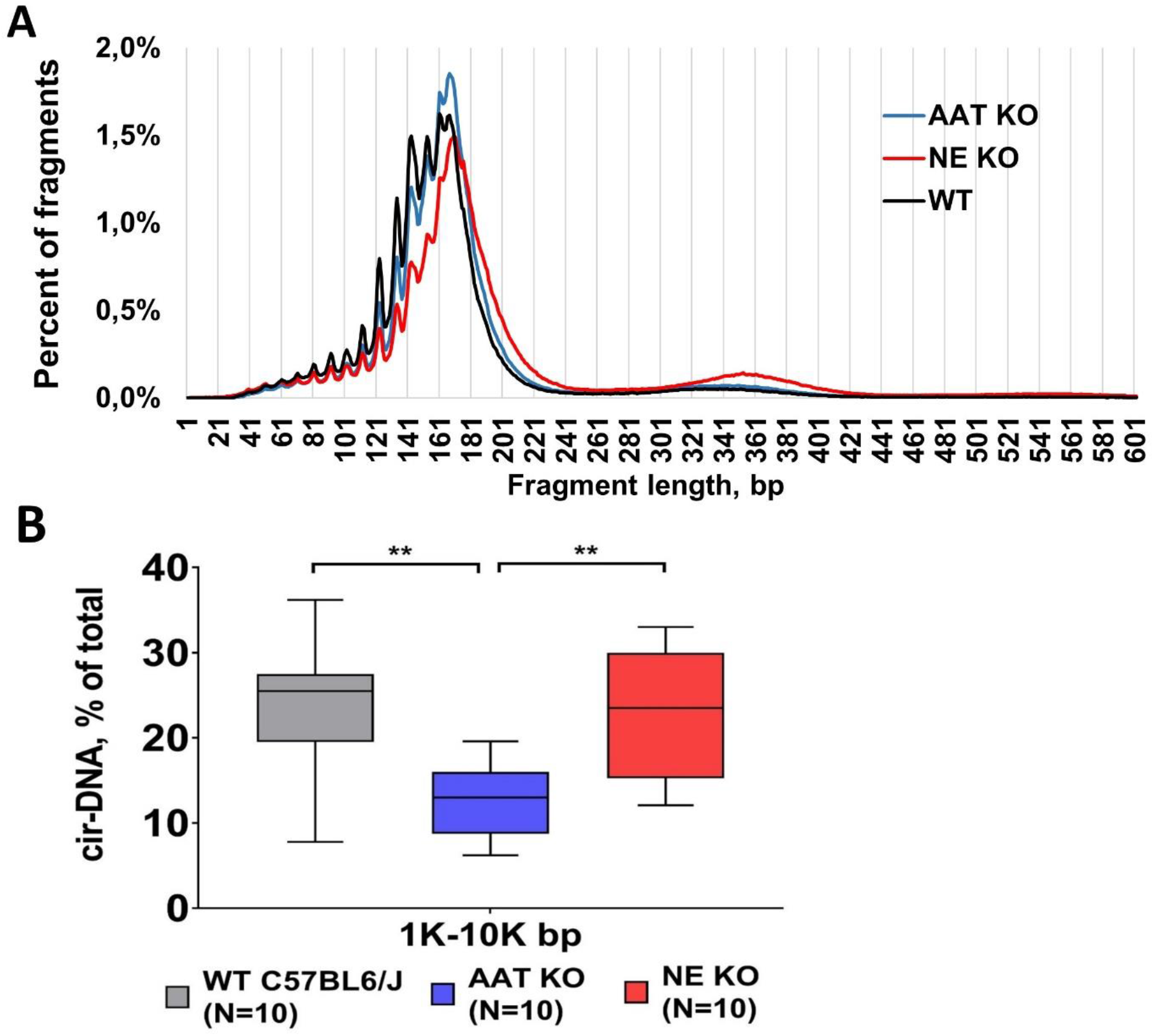
*In vivo* study on the role of the NE in cirDNA fragmentome. (A) Illustrative size profiles of cirDNA from plasma of wild type mice (WT), neutrophil elastase knockout mice (NE KO) and mice knockout for alpha-1 antitrypsin (AAT KO), as determined by sWGS. (B) Analysis of long fragment cirDNA fraction in plasma of WT, NE KO and AAT KO, as determined by capillary electrophoresis (Fragment Analyser, Agilent).

Analysis of cirDNA fractions by capillary electrophoresis revealed a significantly decreased fraction of long DNA fragments in AAT KO mice: the fraction of 1k-10k bp fragments was 2-fold higher in WT and NE KO mice than in AAT KO mice (Fig. 6B).

### NET markers and cirDNA association in plasma in several disorders

Plasma NE, MPO and cirDNA concentrations were significantly elevated in the SLE, mCRC and COVID-19 patients compared to HI (Fig. 7A). The highest values of NET markers were found in the plasma of severe COVID-19 patients, which showed significant differences from HI in the analysis of NE (45.3 ng/ml vs 12.88 ng/ml, p<0.000001), MPO (90.85 ng/ml vs 11.91 ng/ml, p<0.000001) and cirDNA (116 ng/ml vs 5.76 ng/ml, p<0.000001). There was no correlation between cir-nDNA and NE/MPO concentration in HI (Fig. 6B), while they positively correlated in SLE, mCRC and COVID-19 patients (Fig. 7C, D, E). The highest positive correlations of cirDNA with NE and MPO were found in the plasma of SLE patients (r=0.85 and r=0.84, respectively) (Fig. 7E).

**Fig. 7:**
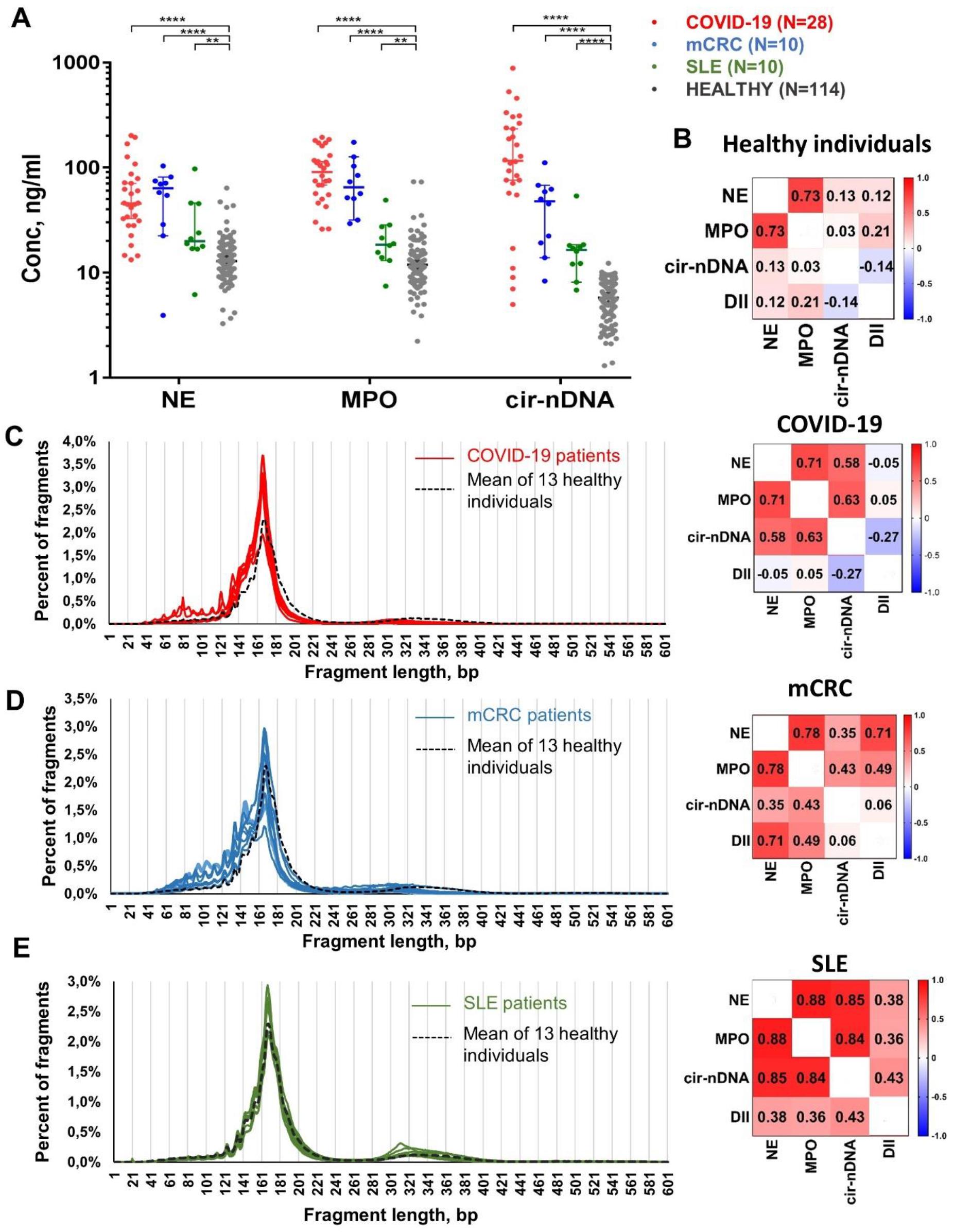
NETs’ association with several disorders. (A) Comparison of cirDNA, MPO and NE concentrations (ng/mL of plasma) in COVID-19 patients (n=28), SLE patients (n=10), mCRC patients (n=10) and healthy individuals (HI) (n=114). Lines represent median with 95% CI. Mann-Whitney test was performed to compare values for cirDNA, MPO and NE in mCRC patients and in HI. A probability of less than 0.05 was considered statistically significant; **p<0.01, ****p< 0.0001. Each dot represents the values of a single patient or a single healthy individual. CirDNA: circulating cell-free DNA; MPO: myeloperoxidase; NE: neutrophil elastase. (B) Correlation matrix of MPO and NE concentrations (ng/mL plasma) with cirDNA parameters in healthy individuals (n=114). Heatmap manifests the strength of the relationship by Pearson’s correlation analysis. Cir-nDNA: circulating cell-free DNA of nuclear origin; MPO: myeloperoxidase; NE: neutrophil elastase; DII: DNA integrity index; cir-mtDNA: circulating cell-free DNA of mitochondrial origin; MNR: ratio of mitochondrial to nuclear circulating DNA concentration. Correlation matrix of MPO and NE concentrations (ng/mL plasma) with cirDNA parameters and comparison of the cfDNA size profile determined by sWGS of the healthy individual mean (black dashed line) and (A) COVID-19 patients, (B) metastatic colorectal cancer (mCRC) patients, and (C) systemic lupus erythematosus (SLE) patients.

As determined by sWGS, the cirDNA size profiles of SLE, mCRC, and COVID-19 patients was somewhat similar to those of HI, which clearly points to mononucleosomes as the predominant structures from which DNA obtains a degree of stability. However, the size profiles of cirDNA from COVID-19 and mCRC patients show a clear shift to shorter fragments, as compared to HI. The di-N associated DNA fragment population from healthy plasma peaked at 332 bp, while it did so at ∼310-320 bp for SLE, mCRC, and COVID-19 patient plasma. The size profile of SLE patients was relatively homogenous and similar to that of HI, except for its di-N peak.

## Discussion

Sufficient evidence now exists regarding NET as one (previously unappreciated) source of cirDNA to indicate their strong potential as a source of biomarkers for various inflammatory disorders and other pathologies. However, no study to date has directly proven that either chromatin or NET DNA fibers degrade into mono-N DNA which could be released into the blood circulation without implying cell death.

### The gHMW DNA can be degraded in serum into mononucleosomes

For more than two decades, the prominence of mononucleosomes in cirDNA extract encouraged the postulation that apoptosis was the main mechanism of release into blood circulation [4,7,39]. Previous efforts to study genomic DNA degradation, however, offered conflicting conclusions regarding the role of apoptosis and apoptotic nucleases in internucleosomal DNA fragmentation and the production of mononucleosomes. On the one hand, certain studies report that cirDNA originates in the apoptotic DNA fragments generated by caspase-activated DNase (CAD), because the cfDNA fragment sizes generally approximate to a mononucleosomal size [40]. Koyama et al. [14] propose that cirDNA can originate from both apoptotic and necrotic cells, and that CAD, DNase γ and DNase I are responsible for the generation of DNA fragments. On the other hand, a different study showed that apoptosis in CAD-deficient mice caused DNA fragmentation equivalent to that in WT mice; thus concluding that apoptosis-associated CAD is therefore not essential for DNA degradation *in vivo*, they suggested that phagocytes are rather as the main actor in DNA fragmentation [41]. Finally, yet another study in mice has shown that, in apoptosis, CAD is insufficient for degrading chromatin, and requires DNase1L3 to degrade it into mononucleosomes [42]. Given these contradictory data and the uncertain role of apoptosis in DNA fragmentation in cell culture or *in vivo* in mice, we preferred to focus on the general ability of human blood to fragment DNA regardless of the mechanisms of cell death [7]. Our current study first investigated DNA degradation by adding gHMW DNA from a cell lysate to the serum or plasma of a HI under physiological conditions meaning that no cells (and thus no apoptosis or any other cell death events) could be involved in DNA degradation. Our study design implies the use of gHMW DNA and blood donors of opposite genders, and exploring the gHMW DNA topology under degradation by examining the *SRY* sequence. An accumulation of DNA fragments associated with mononucleosomes showed that, in the absence of apoptosis, exposure to serum nucleases alone is sufficient for the effective degradation of gHMW DNA down to mono-N DNA fragments. Data also confirmed mono-N as the most stabilized structure in blood with which cirDNA can associate [31,43].

In previous studies, we and other groups have scrutinized the size distribution of cirDNA in the plasma of HI and cancer patients [31,43]. Amongst those findings, a sWGS assay revealed the predominance of chromatosome/mononucleosome-associated cirDNA fragments and the presence in the plasma of HI and cancer patients of a minor population corresponding to di-nucleosome-associated cirDNA. A combination of double- and single-strand DNA library preparations (DSP and SSP) and nested qPCR (N-qPCR) enabled the estimation of the wide size distribution of cirDNA fragments inserted in mono-nucleosomes, di-nucleosomes, and chromatin of higher molecular size (>1000 bp) as 67.5 - 80%, 9.4 - 11.5%, and 8.5 - 21.0%, respectively [31]. We found a striking reproducibility of the cirDNA size profiles in HI [31], in that the maximal size and frequency of the major peak and all subpeaks varied by merely 1 or 2 bp and by 2 - 20%, respectively.

In the present study, we observed a characteristic chromatin pattern after gHMW DNA degradation by serum nucleases, in which the most prominent peak is of mono-N cfDNA, and the less prominent peak of di-N cfDNA (Fig. 2B) as observed for cirDNA *in vivo*. We observed a very rapid degradation and the depletion of long DNA fragments resulted in the formation of oligonucleosomes, making them the main substrate for DNase attacks, and leading to the subsequent accumulation of mono-N and di-N (after 2H incubation). This also led to the accumulation of short DNA fragments in the 40 - 160 bp range, indicating the dynamic action of serum nucleases on the internucleosomal DNA, the DNA linked to the histone H1, and the 14 DNA base pairs exposed at the surface of the nucleosome core particle at a ∼10 bp periodicity [31,43,44].

As determined by sWGS, the analysis of the DNA fragment fractions corresponding to mono-N, di-N and tri-N confirms these conclusions, and adds to our quantitative assessment of the kinetics of gHMW DNA degradation in serum. When comparing the proportions of mono-N, di-N or tri-N DNA, data revealed the higher resistance of mononucleosomes to DNase attacks, as compared to oligonucleosomes. We therefore conclude that in normal physiological conditions blood might be capable of degrading gHMW DNA, and especially nicking and degrading the internucleosomal DNA; this leads to the accumulation of mononucleosomes/chromatasomes.

### Activated neutrophils produce HMW DNA fragments ranging from 1500 – 30 000 bp

Although there are several reports based on the visualization of NET *in vitro* by fluorescent microscopy, which demonstrates that NET consist of the DNA fibers, no studies have analyzed the structure and length of DNA released by activated neutrophils and its association with cirDNA production.

In our present study, we describe the accumulation of HMW DNA fragments and the increase of NET markers in the supernatant of *ex vivo* activated neutrophils. This would indicate the formation of these DNA fibers decorated with NET-specific proteins, such as NE and MPO. Our data confirms these results by revealing the significant correlations of the DNA markers (based on size and amount) with MPO and NE. These results show the association of extracellular DNA with NET, and suggest that NET are the source of the dynamic degradation undergone by HMW DNA in blood.

The MNR negatively correlated with the concentration of NE and MPO, and decreased with the activation of the neutrophils (Fig. 3C and Additional file 1: Fig. S4). In previous studies, we have already described the decrease of MNR [26,35,45] and the increase of NE and MPO in cancer plasma, as compared to healthy individuals [26]. We believe this to be linked to the activation of neutrophils and the release of NET, and could be explained by the structural differences between nuclear and mitochondrial DNA. The release of mitochondrial DNA (mtDNA) by activated neutrophils has been detected in several studies [46,47]. Being devoid of an association with histones (and the protection which results from that), mtDNA is more subject to fragmentation than nuclear DNA, when not inside the circulating mitochondria [48]. We can therefore suggest that the rapid and massive production of both nuclear and mtDNA, combined with the faster degradation of mtDNA compared to the relatively stable nDNA (protected by the mononucleosome structure), leads to an imbalance in the proportions of cir-nDNA and cir-mtDNA; and thus to a decrease of MNR.

### The in vitro degradation in serum of the long DNA fragments originating from ex vivo generated NET leads to the accumulation of the mononucleosome-associated fraction of cfDNA

qPCR quantification revealed the progressive decrease of the total DNA and the long DNA fragment concentrations (Fig. 4A), as well as the synchronous decrease of the DII (Additional file 1: Fig. S5), confirming our postulate that DNA from NET could be degraded by serum nucleases down to mono-N fragments. Analysis of the same DNA extracts by capillary electrophoresis demonstrated the shortening and dynamic reduction of the 1 - 30 kb DNA fraction during incubation in serum (Fig. 4B). That observation agrees with the qPCR results. The analysis of DNA by capillary electrophoresis does not allow us to distinguish male and female DNA using the sequence of the Y chromosome and therefore to directly quantify their respective amount. However, given the 30-fold excess of DNA from NET (0.36 ng/μL) compared to cirDNA from serum (0.012 ng/μL), we can assume that more than 95% of the DNA fragments detected by capillary electrophoresis originate from NET.

The NET production was associated with the accumulation of di-N and tri-N DNA. Conversely, in the course of the degradation of NET DNA in serum we observed an inversion of all those proportions, with the progressive disappearance of di-N and tri-N associated DNA fragments, and the accumulation of mono-N DNA fragments (Fig. 4D). These results confirm our hypothesis that, when expelled to the extracellular milieu (in this case, the serum), the long fragments of DNA originating from NET are predominantly degraded to mononucleosomes, given that (of all NET byproducts) these constitute the most stabilized structure with which cirDNA can associate.

The rate of DNA degradation of NET and gHMW DNA can in no way be compared to previous observations about the half-life of cirDNA in the circulation, where pharmacological aspects such as clearance, mononucleosome phagocytosis or metabolic pathways are heavily involved. Our main objective is to show that the degradation of NET or chromatin leads to mono-N cirDNA. In physiological conditions, detectable cirDNA results from the balance between extracellular DNA release and elimination, as above mentioned. Extracellular release may result from various mechanisms. Of these, NET could be a significant source, along with other mechanisms such as apoptosis, necrosis, active cellular release or extracellular traps generated from other blood cell types [25].

Chromatin organization along the genome is not random, and may be specific to cell gene regulation and cell types [49,50]. CirDNA cell-of-origin was recently investigated using WGS and fragmentomics. Analytical signals from the cirDNA fragmentome include non-random fragmentation pattern, transcription factors occupancy, GC contents, and end motif pattern, and by extension methylation. Studies from Y. Dor’s team based on cirDNA methylome analysis [51,52] appear to indicate that the main cirDNA cells-of-origin are megakaryocytic and lymphocytic (especially neutrophilic). More recent reports confirmed that among a great variety of cells-of-origin, the neutrophil origin appears to be one of the most important [53]. Interestingly, this report and others also found that the cell-of-origin proportion may vary when comparing plasma of HI and non-HI, particularly cancer patients. In addition, cirDNA fragmentation analysis in open chromatin regions [34] and nucleosome occupancy also informs about tissue of origin. Thus, Shendure’s milestone study [44] revealed that lymphoid or myeloid origins have the largest proportions consistent with hematopoietic cells as the dominant source of cirDNA in HI, in contrast to cancer patients. This opens up a wide range of opportunities for diagnosis and the interrogation of physiological and pathological processes [51,54].

### NE affects the cirDNA associated structure in vivo by enhancing DNA fragmentation

NE was found to be involved in DNA decondensation in the nuclei of activated neutrophils [27]. It could therefore be suggested that NE contributes to DNA fragmentation. In our *in vivo* study of cirDNA deriving from the plasma of WT, NE KO and AAT KO mice, we demonstrated that the cirDNA size profiles revealed by sWGS differ according to NE activity. The absence of NE (in NE KO mice) was associated with the increase of the di-N associated cirDNA fragments and a significant increase of the average fragment size, which would indicate impaired DNA fragmentation, as compared to WT mice. The cirDNA fragment size profile of WT and AAT KO mice were visually similar, despite the increased activity of NE in the AAT KO mice. Capillary electrophoresis analysis revealed a decrease in the 1k - 10k bp cirDNA fragment fraction in AAT KO mice, suggesting that NE facilitates or participates in the degradation of DNA fragments of sizes equivalent to those generated from *ex vivo* PMA stimulated neutrophils. Taken together, these results permit the conclusion that NE affects the cirDNA structure and, consequently, the fragmentation features of cirDNA which originates from NET.

### NE and MPO synergistically catabolize NET

The mechanism of cirDNA generation, in particular its enzymatic activity, remains insufficiently understood. Theoretically, the catabolism of DNA which leads to cirDNA production may be enabled by nucleases either intracellularly (DNase1L1, DNA fragmentation factor B or DFFB, DFFB/CAD and endonuclease G) or extracellularly (DNase1 and DNase1L3 or DNase γ). The paradigm in which, historically, this mechanism has been understood involved mainly apoptosis as well as necrosis [4,7]. This is due to initial observations of laddered patterns of oligonucleosomal-size pieces (being the hallmarks of canonical apoptosis) by gel electrophoresis of cirDNA extract, and due also to the more recent characterization of the mononucleosome structure, principally by sequencing and fragmentomics [31,43,44,55]. The main nuclease responsible for the internucleosomal double strand (ds) cleavage of genomic DNA is DFFB/CAD; it may also be responsible, however, for single-strand breaks (ssB) during apoptosis [56]. Watanabe et al observed that DNase1L3 cooperated with caspase-activated DNase (CAD) in the case of anti-Fas mediated hepatocyte apoptosis, while DNase1L3 was responsible only for the generation of cirDNA in acetaminophen-induced hepatocyte necrosis. However, they were unable to detect cirDNA in CAD-/- DNase1L3-/- mice, indicating that DNase1 is not sufficient for the generation of cirDNA following apoptosis [42]. This supported the observations made by D. Lo’s group [57,58], who reported similar cirDNA levels in both DNase1 -/- and WT mice in steady state condition. Although the mechanism of DNase activity is not fully understood, it shows correlation with inflammation; for instance, its induction mechanism may be associated with macrophage activation [59]. Collating various observations [14,42,58,60] provide clues that CAD plays a key role in the initiation of cell death, by producing dsDNA breaks intracellularly; while DNase1L3 and DNase1 (both induced by apoptosis and necrosis) pursue chromatin catabolism extracellularly, such as in blood circulation. DNase1 is expressed by non-hematopoietic tissues and preferentially cleaves protein-free DNA [61,62]. DNase1L3 (also known as DNase gamma) is secreted by immune cells, and targets DNA-protein complexes such as nucleosomes [59,62]. Note, several reports have postulated the role of active release in the production of DNA associated to extracellular vesicles [63]. The literature showed discrepancies about DNA being incorporated onto or into microvesicles. DNA encapsulated in vesicles would be less sensitive to nuclease attack, potentially modifying DNA fragmentation rate and process. Conversely, it was demonstrated that nuclease DNA1L3 could degrade DNA in microvesicles [59]. Regardless of these considerations, once released from vesicles its fragmentation process should be similar to that of cirDNA not associated with extracellular vesicles . Over the last five years, we [3] and several other authors [42,64,65] have speculated about the role of NETosis in the generation of cirDNA. There are conflicting observations as to the nature of the nuclease-derived enzymatic activity involved in the degradation of NET to cirDNA. Both DNase1 and DNase1L3 have been shown to degrade NET in blood circulation [65,66]. Jimenez et al. [65] showed that the toleration of chronic neutrophilia (as well as the prevention of vascular occlusion by NET clots during that process) requires both enzymes.

Data suggests that DNA degradation is increased when NE and/or MPO is added to serum nucleases, and that DNA degradation is increased to a much larger extent in plasma when nuclease activity is greatly inhibited. Data strikingly showed that, in serum and in plasma, the DNA degradation rate due to NE was nearly equivalent, as it was (to a higher extent) due to MPO activities, irrespective of nuclease inhibition by EDTA as chelating agent. Note, NE activity was found not dependent of Ca and Mg ions and determined in plasma [67]. Whereas MPO is dependant of Ca ions, its activity was still present in plasma [68]. This was the first direct observation of NE and MPO DNA degradation activity. Overall, our study consequently suggests that NE and MPO contribute to DNA catabolism. In addition, our data clearly demonstrated, for the first time, that NE and MPO contribute to and favor the DNA degradation process by releasing oligo-nucleosomes, maily mono-N from NET. In addition, we suggest that, following neutrophil stimulation principally by oxidative explosion and its translocation to the nucleus, NE is continuously released from extracellular NET-associated granules, and continuously decondenses the chromatin. It has been established that histones are cleaved by NE derived from granulation, and are citrullinated by peptidyl arginine deiminase 4 (PAD4) during intracellular NET formation. In the initial phase of this process, NE degrades the linker protein H1 and then the core histone proteins, in a process which is known to be enhanced by MPO [27]. MPO produces oxidizing agents such as hypochlorous acid (HOCl) from hydrogen peroxide (H2O2) and chloride anion (Cl−). Thus, we might postulate another possible hypothesis: oxidizers damage DNA, thus making it more susceptible to degradation by nucleases [69]. Drugs producing oxidizers might therefore be conceived as NET inhibitors, in pathologies where NET formation is uncontrolled [25].

In addition, our study first revealed that the synergistic association of NE and MPO contributes to the degradation of *in vitro* gHMW DNA, leading to the release of oligo- and principally mono-nucleosomes in serum containing medium. However, our study is limited since it can not fully establish whether NE/MPO contribute to NETs degradation either directly (such as having nuclease activity) or indirectly (such as chromatin unpacking, facilitating nuclease attack). Nonetheless, taking these observations as a whole, we suggest that the combined action of NE and MPO may have a dual role, firstly in intracellularly initiating NET formation, and secondly in participating in NET degradation in the extracellular milieu (i.e., blood), which would point to a potential NET autocatabolism.

Conversely, other NET constituents may be involved in protecting DNA from nucleases such as the antimicrobial peptide LL-37 [70]. Alternatively, thrombin may bind to NET, thus conferring mutual protection against nuclease and protease degradation [71], illustrating the complex interplay between coagulation and NET formation [72].

Note, we found low but significant levels of both enzymes in the circulation of healthy individuals. We believe that the homeostasis of both enzymes and of nucleases such as DNase1L3 or DNase1 is critical in controlling NET formation on the one hand, and cirDNA plasma concentration on the other. We speculate that genetic or epigenetic alterations impacting the expression of these entities may mitigate or exacerbate both the damage-associated molecular patterns (DAMPs) of cirDNA [3,73] and NET formation, leading to chronic or acute disorders.

### NET association with several disorders

NET formation is an efficient, innate response strategy for counteracting intrusive micro-organisms. However, exacerbated NET formation in the host can be detrimental, on account of the toxicity of its exposed compounds to endothelial cells and to parenchymal tissue [54], whose pathological consequences can include thrombosis and fibrinolysis disorders. For instance, NET components (DNA, histones, and granule proteins such as MPO or NE) may trigger an inflammatory process. Thus, the release of NET byproducts as a result of dysregulated NET formation, therefore, can be implicated in both autoimmune and non-autoimmune diseases [74–76].

Intense investigation has been done on the deleterious role of NET in autoimmune diseases, particularly in the pathogenesis of systemic lupus erythematosus (SLE). The dysregulation of NET formation has been repeatedly demonstrated in SLE [77,78]; patients with SLE, notably, have increased levels of NET in their tissues and circulation. Thus, NET lead to the exposition of many autoantigens within the extracellular space, in particular double strand DNA, and MPO. As a result, autoimmune complexes containing nucleic acids associated with various proteins (including antibodies, the chromatin-associated protein HMGB1, the antimicrobial peptide LL39, and ribonucleoproteins) are found in the sera of SLE patients [78]. Evidence has also been presented that the ability to degrade NET is impaired in patients with SLE [79], leading to a continuous release of interferon. Since this key cytokine in the SLE pathophysiology is also responsible for sensitizing neutrophils to NET formation, the overall result is the establishment of an amplifier loop which (with other factors) enables the autoimmune reaction to be sustained, and the disease to progress. Given all of the above, NET byproducts are now used as biomarkers for SLE clinical diagnostics [80]. In our study, we found a strong association of cirDNA with NE and MPO, as well as a negative correlation of MNR with NET-related biomarkers in the plasma of SLE patients [81]. These results are similar to those found in the supernatant of *ex vivo* activated neutrophils (Fig. 3), suggesting NETs’ involvement in the pathogenesis of SLE.

Several recent reports have also described the involvement of NET formation in cancer progression [82–86]. Some authors even postulate that NET are a seedbed for metastasis [86]. We recently demonstrated the association of MPO and NE concentrations with the amounts of cirDNA and anti-cardiolipin auto-antibodies in a large cohort (N=217) of newly diagnosed metastatic colorectal cancer (mCRC) patients [26]. In this current study, we clearly confirmed this observation in a small group of mCRC patients, in whom we found a statistically significant higher level of NE, MPO and cirDNA plasma concentration, as compared to healthy individuals.

We are among the first to indicate that NET and NET byproducts play a key role in COVID-19 pathogenesis [25,76,87]. In this regard, notably, we have shown strong analogies between the multiple biological and pathological features, risk factors, and general clinical conditions observed in COVID-19 with several other disorders in which the uncontrolled formation of NET and their by-products is implicated [25,76,87]. We also hypothesized that just such a deregulated NET formation is critically involved in the progression of COVID-19 under an amplifier loop, in a massive, uncontrolled inflammatory process [76]. Zuo et al [88] were the first to experimentally show that cirDNA as well as NET protein byproducts are associated with COVID-19, and might be potential COVID-19 markers. Like a number of other authors, we have here confirmed that NE, MPO and cirDNA concentrations in the plasma of patients with severe COVID-19 are significantly higher than in a large cohort of healthy individuals. We are now exploring the clinical validation of NET biomarkers in an ongoing clinical study which includes 160 patients.

Altogether, we found a significant increase in both NET proteins and cirDNA concentration in the plasma of SLE, mCRC and COVID-19 patients (Fig. 7). Both NE and MPO with cirDNA markers (DII, cir-nDNA) showed strong positive correlations in patients with these three inflammatory diseases. These results confirm our *in vitro* and *ex vivo* experiments, adding further weight to the case for NET being one of the source of cirDNA. We also found a constant and strong negative correlation of MNR with NET biomarkers, suggesting that the release and degradation rate of mitochondrial DNA could be associated with NET production, resulting in a decrease of the MNR in NET-associated disorders.

The topological features and extracellular mechanisms of the degradation of cir-mtDNA appear to be different from those of cir-nDNA, and are currently under intense investigation in our laboratory [25,48]. To these observations must be added those of our current study, which is the first to demonstrate that the cirDNA size profile of subjects suffering from SLE, mCRC and COVID-19 has features distinct from that of healthy individuals, notably a cirDNA fragmentation specifically induced by a high level of NET formation. Overall, the size profiles of COVID-19 and mCRC patients, and to a lesser extent SLE patients, demonstrate a subtle but significant shift towards short DNA fragments, all of which indicates increased fragmentation of cirDNA. Note, the study is somewhat limited by the number of subjects, especially as compared to a clinical assay. The number of subjects nonetheless allowed us to observe significant statistical differences between groups and to draw preliminary observations. These will have to be confirmed with a larger number of patients to obtain more solid conclusions as to the impact of pathological NET dysregulation. Our study on quantitative values, association assays, and cirDNA size profile showed significant differences in patients versus HI, but remains exploratory and must be confirmed in larger patient cohorts. Overall, this exploratory work on the plasma of patients with diseases implying high levels of both NET formation and cirDNA (as compared to the plasma of HI) suggests (i) a specific and high association of cirDNA amount with NET protein markers; and, (ii) a modification of the fragmentome of cirDNA from these patients, in whom NET may therefore be a significant source of cirDNA. This may imply that the proportion of NET as a cirDNA source is higher in these diseases, or at least that in these patients there is a difference in the respective proportion of the various mechanisms of cirDNA release mentioned above.

## Conclusions

The detection of cirDNA in blood is characteristically associated with the protective effect of mononucleosomes, which determines its fragmentation. The co-occurring observation that mononucleosomes clearly accumulate in cfDNA extracts following incubation of gHMW DNA in serum, or in plasma cirDNA extracts from mice or human subjects, in combination with the observation that cultured activated neutrophil produced NET and high molecular weight DNA that are further degraded mainly to mononucleosomes when incubated in cell culture medium where apoptosis cannot occur, strongly suggests that in a physiological setting NET may produce cirDNA as a byproduct, independent of apoptosis. The contribution of this process to cirDNA levels compared to other known mechanisms of release, such as apoptosis or necrosis, needs to be evaluated in healthy individuals, and may have diagnostic potential with regard to diseases with NETopathies, such as some inflammatory diseases. Our work thus describes the previously unknown mechanistic bridge linking NET and cirDNA, and establishes a new paradigm of the mechanism of cirDNA release in both normal and pathological conditions. In addition, our observations establish a link between immune response and cirDNA, thus enlarging the scope of the diagnostic potential offered by cirDNA, and giving a better understanding of the pathogenesis of inflammatory diseases.

## Abbreviations

NET: neutrophil extracellular traps;
gHMW DNA: genomic high molecular weight DNA;
NE: neutrophil elastatse;
MPO: myeloperoxidase;
mono-N: mononucleosome;
di-N: dinucleosome;
tri-n: trinucleosome;
cirDNA: circulating DNA;
cir-nDNA: circulating DNA of nuclear origin;
cir-mtDNA: circulating DNA of mitochondrial origin;
LPS: lipopolysaccharides;
PMA: Phorbol 12-myristate 13- acetate;
qPCR: quantitative polymerase chain reaction;
sWGS: shallow whole genome sequencing;
SLE: systemic lupus erythematosus;
mCRC: metastatic colorectal cancer;
HI: healthy individuals.

## Declarations

### Ethics approval and consent to participate

Plasma samples from 28 patients with severe COVID-19 (at the entrance of Intensive Care Unit) were provided by the CHU hospital of Montpellier (Centre Hospitalier Universitaire de Montpellier, France); approval number assigned by the IRB : IRB-MTP_2021_01_202100715. We also included 10 mCRC patients from the screening procedure of the ongoing UCGI 28 PANIRINOX study (NCT02980510/EudraCT n°2016-001490-33). Plasma samples from 10 female patients with systemic lupus erythematosus (SLE) were provided by Dr. P. Simon from the Department of Sports Medicine, Prevention and Rehabilitation of Johannes Gutenberg University (Mainz, Germany). The study was approved by the Human Ethics Committee of Rhineland-Palatinate, Germany (number 2018-13039) and conformed to the standards of the Declaration of Helsinki of the World Medical Association and was registered under ClinicalTrials.gov Identifier: NCT03942718. We also analyzed 114 healthy individuals (HI, 59 men and 55 women) from the Etablissement Français du Sang (EFS), which is Montpellier’s blood transfusion center (Convention EFS-PM N° 21PLER2015-0013).

### Consent for publication

Not applicable.

### Availability of data and materials

The sWGS data described in this article can be freely and openly accessed at Zenodo: https://doi.org/10.5281/zenodo.6949112

### Authors’ contributions

ART and EP designed the study. EP and ART developed the methodology. EP, LM, BP, AK, and AM did the experiments under the supervision of ART. EP did the statistical analyses. EP and ART analyzed the data and prepared the manuscript. SB provided the plasma samples of COVID-19 patients. UM provided the samples of WT, NE KO and AAT KO mice. SCB, JWM, EWIN, and PS provided the plasma samples of SLE patients. All of the authors (EP, LM, BP, AK, AM, TM, SB, UM, SCB, JWM, EWIN, PS and ART) discussed the results and approved the manuscript.

## Acknowledgments

The authors thank the excellent technical assistance of C. Sanchez and F. Frayssinoux (IRCM, Institut de Recherche en Cancérologie de Montpellier, INSERM U1194, Université de Montpellier, Institut régional du Cancer de Montpellier, Montpellier, F-34134298, France); J.P. Cristol, K. Klouche, P. Fessler, C. Roubille, M. Berger (CHU Montpellier, France); M. Ychou (ICM, Institute of Cancerology of Montpellier); Simone C. Boedecker (University Medical Center Mainz, Germany) and Cormac Mc Carthy (Mc Carthy Consultant, Montpellier) for English editing (financial compensation). D. Tousch (University of Montpellier, France) for reading the manuscript. Figure 1 was created with BioRender.com

## References

1. Bronkhorst AJ, Ungerer V, Diehl F, Anker P, Dor Y, Fleischhacker M, et al. Towards systematic nomenclature for cell-free DNA. Hum Genet [Internet]. 2020 [cited 2020 Nov 6]; Available from: http://link.springer.com/10.1007/s00439-020-02227-2

2. Lo YMD. Non-invasive prenatal diagnosis by massively parallel sequencing of maternal plasma DNA. Open Biol [Internet]. 2012 [cited 2020 May 15];2. Available from: https://www.ncbi.nlm.nih.gov/pmc/articles/PMC3390796/

3. Thierry AR, El Messaoudi S, Gahan PB, Anker P, Stroun M. Origins, structures, and functions of circulating DNA in oncology. Cancer Metastasis Rev. 2016;35:347–76.

4. Jahr S, Hentze H, Englisch S, Hardt D, Fackelmayer FO, Hesch R-D, et al. DNA Fragments in the Blood Plasma of Cancer Patients: Quantitations and Evidence for Their Origin from Apoptotic and Necrotic Cells. Cancer Res. American Association for Cancer Research; 2001;61:1659–65.

5. Holdenrieder S, Mueller S, Stieber P. Stability of nucleosomal DNA fragments in serum. Clin Chem. 2005;51:1026–9.

6. Deligezer U, Erten N, Akisik EE, Dalay N. Circulating fragmented nucleosomal DNA and caspase-3 mRNA in patients with lymphoma and myeloma. Exp Mol Pathol. 2006;80:72–6.

7. Rostami A, Lambie M, Yu CW, Stambolic V, Waldron JN, Bratman SV. Senescence, Necrosis, and Apoptosis Govern Circulating Cell-free DNA Release Kinetics. Cell Rep. 2020;31:107830.

8. Barra GB, Santa Rita TH, Vasques J de A, Chianca CF, Nery LFA, Costa SSS. EDTA-mediated inhibition of DNases protects circulating cell-free DNA from ex vivo degradation in blood samples. Clinical Biochemistry. 2015;48:976–81.

9. Li LY, Luo X, Wang X. Endonuclease G is an apoptotic DNase when released from mitochondria. Nature. 2001;412:95–9.

10. Liu X, Zou H, Slaughter C, Wang X. DFF, a heterodimeric protein that functions downstream of caspase-3 to trigger DNA fragmentation during apoptosis. Cell. 1997;89:175–84.

11. Parrish J, Li L, Klotz K, Ledwich D, Wang X, Xue D. Mitochondrial endonuclease G is important for apoptosis in C. elegans. Nature. 2001;412:90–4.

12. Mizuta R, Mizuta M, Araki S, Shiokawa D, Tanuma S, Kitamura D. Action of apoptotic endonuclease DNase gamma on naked DNA and chromatin substrates. Biochem Biophys Res Commun. 2006;345:560–7.

13. Mizuta R, Araki S, Furukawa M, Furukawa Y, Ebara S, Shiokawa D, et al. DNase γ is the effector endonuclease for internucleosomal DNA fragmentation in necrosis. PLoS ONE. 2013;8:e80223.

14. Koyama R, Arai T, Kijima M, Sato S, Miura S, Yuasa M, et al. DNase γ, DNase I and caspase-activated DNase cooperate to degrade dead cells. Genes Cells. 2016;21:1150–63.

15. Herriott RM, Connolly JH, Gupta S. Blood Nucleases and Infectious Viral Nucleic Acids. Nature. Nature Publishing Group; 1961;189:817–20.

16. Tamkovich SN, Cherepanova AV, Kolesnikova EV, Rykova EY, Pyshnyi DV, Vlassov VV, et al. Circulating DNA and DNase activity in human blood. Ann N Y Acad Sci. 2006;1075:191–6.

17. Brinkmann V, Reichard U, Goosmann C, Fauler B, Uhlemann Y, Weiss DS, et al. Neutrophil extracellular traps kill bacteria. Science. 2004;303:1532–5.

18. Pilsczek FH, Salina D, Poon KKH, Fahey C, Yipp BG, Sibley CD, et al. A novel mechanism of rapid nuclear neutrophil extracellular trap formation in response to Staphylococcus aureus. J Immunol. 2010;185:7413–25.

19. Hakkim A, Fürnrohr BG, Amann K, Laube B, Abed UA, Brinkmann V, et al. Impairment of neutrophil extracellular trap degradation is associated with lupus nephritis. PNAS. National Academy of Sciences; 2010;107:9813–8.

20. Kaplan MJ, Radic M. Neutrophil extracellular traps: double-edged swords of innate immunity. J Immunol. 2012;189:2689–95.

21. Luo L, Zhang S, Wang Y, Rahman M, Syk I, Zhang E, et al. Proinflammatory role of neutrophil extracellular traps in abdominal sepsis. Am J Physiol Lung Cell Mol Physiol. 2014;307:L586–596.

22. Fuchs TA, Kremer Hovinga JA, Schatzberg D, Wagner DD, Lämmle B. Circulating DNA and myeloperoxidase indicate disease activity in patients with thrombotic microangiopathies. Blood. 2012;120:1157–64.

23. Daniel C, Leppkes M, Muñoz LE, Schley G, Schett G, Herrmann M. Extracellular DNA traps in inflammation, injury and healing. Nat Rev Nephrol. 2019;15:559–75.

24. Pastor B, Abraham J-D, Pisareva E, Sanchez C, Kudriavstev A, Tanos R, et al. Association of the Neutrophil Extracellular Traps Formation With the Production of Circulating Cell-Free DNA and Anti-Cardiolipin Autoantibody in Patients With a Metastatic Colorectal Cancer [Internet]. Rochester, NY: Social Science Research Network; 2021 Aug. Report No.: ID 3912217. Available from: https://papers.ssrn.com/abstract=3912217

25. Thierry AR, Roch B. Neutrophil Extracellular Traps and By-Products Play a Key Role in COVID-19: Pathogenesis, Risk Factors, and Therapy. J Clin Med. 2020;9:E2942.

26. Pastor B, Abraham J-D, Pisareva E, Sanchez C, Kudriavstev A, Tanos R, et al. Association of neutrophil extracellular traps with the production of circulating DNA in patients with colorectal cancer. iScience. 2022;25:103826.

27. Papayannopoulos V, Metzler KD, Hakkim A, Zychlinsky A. Neutrophil elastase and myeloperoxidase regulate the formation of neutrophil extracellular traps. J Cell Biol. 2010;191:677–91.

28. Thierry AR. Circulating DNA fragmentomics and cancer screening. Cell Genomics. 2022;

29. Mouliere F, Robert B, Arnau Peyrotte E, Del Rio M, Ychou M, Molina F, et al. High fragmentation characterizes tumour-derived circulating DNA. PLoS ONE. 2011;6:e23418.

30. Thierry A, Molina F. Analytical Methods for Cell Free Nucleic Acids and Applications [Internet]. 2012 [cited 2021 Dec 16]. Available from: https://patentscope.wipo.int/search/en/detail.jsf?docId=WO2012028746

31. Sanchez C, Roch B, Mazard T, Blache P, Dache ZAA, Pastor B, et al. Circulating nuclear DNA structural features, origins, and complete size profile revealed by fragmentomics. JCI Insight. 2021;6:144561.

32. Cristiano S, Leal A, Phallen J, Fiksel J, Adleff V, Bruhm DC, et al. Genome-wide cell-free DNA fragmentation in patients with cancer. Nature. 2019;570:385–9.

33. Mouliere F, Chandrananda D, Piskorz AM, Moore EK, Morris J, Ahlborn LB, et al. Enhanced detection of circulating tumor DNA by fragment size analysis. Sci Transl Med. 2018;10.

34. Sun K, Jiang P, Cheng SH, Cheng THT, Wong J, Wong VWS, et al. Orientation-aware plasma cell-free DNA fragmentation analysis in open chromatin regions informs tissue of origin. Genome Res. 2019;29:418–27.

35. Tanos R, Tosato G, Otandault A, Al Amir Dache Z, Pique Lasorsa L, Tousch G, et al. Machine Learning-Assisted Evaluation of Circulating DNA Quantitative Analysis for Cancer Screening. Adv Sci (Weinh). 2020;7:2000486.

36. Meddeb R, Pisareva E, Thierry AR. Guidelines for the Preanalytical Conditions for Analyzing Circulating Cell-Free DNA. Clin Chem. 2019;65:623–33.

37. Mouliere F, El Messaoudi S, Gongora C, Guedj A-S, Robert B, Del Rio M, et al. Circulating Cell-Free DNA from Colorectal Cancer Patients May Reveal High KRAS or BRAF Mutation Load. Transl Oncol. 2013;6:319–28.

38. Ostermann L, Maus R, Stolper J, Schütte L, Katsarou K, Tumpara S, et al. Alpha-1 antitrypsin deficiency impairs lung antibacterial immunity in mice. JCI Insight. 2021;6.

39. Grabuschnig S, Bronkhorst AJ, Holdenrieder S, Rosales Rodriguez I, Schliep KP, Schwendenwein D, et al. Putative Origins of Cell-Free DNA in Humans: A Review of Active and Passive Nucleic Acid Release Mechanisms. Int J Mol Sci. 2020;21.

40. Widlak P, Li P, Wang X, Garrard WT. Cleavage Preferences of the Apoptotic Endonuclease DFF40 (Caspase-activated DNase or Nuclease) on Naked DNA and Chromatin Substrates *. Journal of Biological Chemistry. Elsevier; 2000;275:8226–32.

41. McIlroy D, Tanaka M, Sakahira H, Fukuyama H, Suzuki M, Yamamura K, et al. An auxiliary mode of apoptotic DNA fragmentation provided by phagocytes. Genes Dev. 2000;14:549–58.

42. Watanabe T, Takada S, Mizuta R. Cell-free DNA in blood circulation is generated by DNase1L3 and caspase-activated DNase. Biochem Biophys Res Commun. 2019;516:790–5.

43. Sanchez C, Snyder MW, Tanos R, Shendure J, Thierry AR. New insights into structural features and optimal detection of circulating tumor DNA determined by single-strand DNA analysis. NPJ Genom Med. 2018;3:31.

44. Snyder MW, Kircher M, Hill AJ, Daza RM, Shendure J. Cell-free DNA Comprises an In Vivo Nucleosome Footprint that Informs Its Tissues-Of-Origin. Cell. 2016;164:57–68.

45. Meddeb R, Dache ZAA, Thezenas S, Otandault A, Tanos R, Pastor B, et al. Quantifying circulating cell-free DNA in humans. Sci Rep. 2019;9:5220.

46. Yousefi S, Gold JA, Andina N, Lee JJ, Kelly AM, Kozlowski E, et al. Catapult-like release of mitochondrial DNA by eosinophils contributes to antibacterial defense. Nat Med. 2008;14:949–53.

47. Yousefi S, Mihalache C, Kozlowski E, Schmid I, Simon HU. Viable neutrophils release mitochondrial DNA to form neutrophil extracellular traps. Cell Death Differ. 2009;16:1438–44.

48. Al Amir Dache Z, Otandault A, Tanos R, Pastor B, Meddeb R, Sanchez C, et al. Blood contains circulating cell-free respiratory competent mitochondria. FASEB J. 2020;34:3616–30.

49. Roychowdhury T, Abyzov A. Chromatin organization modulates the origin of heritable structural variations in human genome. Nucleic Acids Res. 2019;47:2766–77.

50. Audit B, Zaghloul L, Baker A, Arneodo A, Chen C-L, d’Aubenton-Carafa Y, et al. Megabase replication domains along the human genome: relation to chromatin structure and genome organisation. Subcell Biochem. 2013;61:57–80.

51. Moss J, Magenheim J, Neiman D, Zemmour H, Loyfer N, Korach A, et al. Comprehensive human cell-type methylation atlas reveals origins of circulating cell-free DNA in health and disease. Nat Commun [Internet]. 2018 [cited 2020 Apr 15];9. Available from: https://www.ncbi.nlm.nih.gov/pmc/articles/PMC6265251/

52. Lehmann-Werman R, Neiman D, Zemmour H, Moss J, Magenheim J, Vaknin-Dembinsky A, et al. Identification of tissue-specific cell death using methylation patterns of circulating DNA. Proc Natl Acad Sci U S A. 2016;113:E1826–1834.

53. Sadeh R, Sharkia I, Fialkoff G, Rahat A, Gutin J, Chappleboim A, et al. Author Correction: ChIP-seq of plasma cell-free nucleosomes identifies gene expression programs of the cells of origin. Nat Biotechnol. 2021;39:642.

54. Cheng OZ, Palaniyar N. NET balancing: a problem in inflammatory lung diseases. Front Immunol. 2013;4:1.

55. Ivanov M, Baranova A, Butler T, Spellman P, Mileyko V. Non-random fragmentation patterns in circulating cell-free DNA reflect epigenetic regulation. BMC Genomics. 2015;16 Suppl 13:S1.

56. Iglesias-Guimarais V, Gil-Guiñon E, Sánchez-Osuna M, Casanelles E, García-Belinchón M, Comella JX, et al. Chromatin Collapse during Caspase-dependent Apoptotic Cell Death Requires DNA Fragmentation Factor, 40-kDa Subunit-/Caspase-activated Deoxyribonuclease-mediated 3′-OH Single-strand DNA Breaks *. Journal of Biological Chemistry. Elsevier; 2013;288:9200–15.

57. Cheng THT, Lui KO, Peng XL, Cheng SH, Jiang P, Chan KCA, et al. DNase1 Does Not Appear to Play a Major Role in the Fragmentation of Plasma DNA in a Knockout Mouse Model. Clin Chem. 2018;64:406–8.

58. Serpas L, Chan RWY, Jiang P, Ni M, Sun K, Rashidfarrokhi A, et al. Dnase1l3 deletion causes aberrations in length and end-motif frequencies in plasma DNA. Proc Natl Acad Sci USA. 2019;116:641–9.

59. Sisirak V, Sally B, D’Agati V, Martinez-Ortiz W, Özçakar ZB, David J, et al. Digestion of Chromatin in Apoptotic Cell Microparticles Prevents Autoimmunity. Cell. 2016;166:88–101.

60. Kawane K, Fukuyama H, Yoshida H, Nagase H, Ohsawa Y, Uchiyama Y, et al. Impaired thymic development in mouse embryos deficient in apoptotic DNA degradation. Nat Immunol. 2003;4:138–44.

61. Napirei M, Ricken A, Eulitz D, Knoop H, Mannherz HG. Expression pattern of the deoxyribonuclease 1 gene: lessons from the Dnase1 knockout mouse. Biochem J. 2004;380:929–37.

62. Napirei M, Ludwig S, Mezrhab J, Klöckl T, Mannherz HG. Murine serum nucleases--contrasting effects of plasmin and heparin on the activities of DNase1 and DNase1-like 3 (DNase1l3). FEBS J. 2009;276:1059–73.

63. Cai J, Han Y, Ren H, Chen C, He D, Zhou L, et al. Extracellular vesicle-mediated transfer of donor genomic DNA to recipient cells is a novel mechanism for genetic influence between cells. J Mol Cell Biol. 2013;5:227–38.

64. Beiter T, Fragasso A, Hudemann J, Schild M, Steinacker J, Mooren FC, et al. Neutrophils release extracellular DNA traps in response to exercise. Journal of Applied Physiology. American Physiological Society; 2014;117:325–33.

65. Jiménez-Alcázar M, Rangaswamy C, Panda R, Bitterling J, Simsek YJ, Long AT, et al. Host DNases prevent vascular occlusion by neutrophil extracellular traps. Science. 2017;358:1202–6.

66. Nakayama T, Saitoh H. Tunicamycin-induced neutrophil extracellular trap (NET)-like structures in cultured human myeloid cell lines. Cell Biol Int. 2015;39:355–9.

67. Kunder M, Kutty AM, Lakshmaiah V, Sheela SR. Correlation of Plasma Neutrophil Elastase Activity and Endogenous Protease Inhibitor Levels with the Severity of Pre-eclampsia. J Clin Diagn Res. 2017;11:BC09–BC12.

68. Kothari N, Keshari RS, Bogra J, Kohli M, Abbas H, Malik A, et al. Increased myeloperoxidase enzyme activity in plasma is an indicator of inflammation and onset of sepsis. J Crit Care. 2011;26:435.e1-7.

69. Cooke MS, Evans MD, Dizdaroglu M, Lunec J. Oxidative DNA damage: mechanisms, mutation, and disease. The FASEB Journal. 2003;17:1195–214.

70. Neumann A, Völlger L, Berends ETM, Molhoek EM, Stapels DAC, Midon M, et al. Novel role of the antimicrobial peptide LL-37 in the protection of neutrophil extracellular traps against degradation by bacterial nucleases. J Innate Immun. 2014;6:860–8.

71. Saravanan R, Choong YK, Lim CH, Lim LM, Petrlova J, Schmidtchen A. Cell-Free DNA Promotes Thrombin Autolysis and Generation of Thrombin-Derived C-Terminal Fragments. Front Immunol. 2021;12:593020.

72. Thierry AR. Anti-protease Treatments Targeting Plasmin(ogen) and Neutrophil Elastase May Be Beneficial in Fighting COVID-19. Physiol Rev. 2020;100:1597–8.

73. Denning N-L, Aziz M, Gurien SD, Wang P. DAMPs and NETs in Sepsis. Front Immunol. 2019;10:2536.

74. Barbosa da Cruz D, Helms J, Aquino LR, Stiel L, Cougourdan L, Broussard C, et al. DNA-bound elastase of neutrophil extracellular traps degrades plasminogen, reduces plasmin formation, and decreases fibrinolysis: proof of concept in septic shock plasma. FASEB J. 2019;33:14270–80.

75. Camp JV, Jonsson CB. A Role for Neutrophils in Viral Respiratory Disease. Front Immunol. 2017;8:550.

76. Thierry AR, Roch B. SARS-CoV2 may evade innate immune response, causing uncontrolled neutrophil extracellular traps formation and multi-organ failure. Clin Sci (Lond). 2020;134:1295–300.

77. Boeltz S, Amini P, Anders H-J, Andrade F, Bilyy R, Chatfield S, et al. To NET or not to NET:current opinions and state of the science regarding the formation of neutrophil extracellular traps. Cell Death Differ. 2019;26:395–408.

78. Lamphier M, Zheng W, Latz E, Spyvee M, Hansen H, Rose J, et al. Novel Small Molecule Inhibitors of TLR7 and TLR9: Mechanism of Action and Efficacy In Vivo. Mol Pharmacol. American Society for Pharmacology and Experimental Therapeutics; 2014;85:429–40.

79. Elkon KB. Review: Cell Death, Nucleic Acids, and Immunity: Inflammation Beyond the Grave. Arthritis Rheumatol. 2018;70:805–16.

80. Lee KH, Kronbichler A, Park DD-Y, Park Y, Moon H, Kim H, et al. Neutrophil extracellular traps (NETs) in autoimmune diseases: A comprehensive review. Autoimmun Rev. 2017;16:1160–73.

81. Neuberger EWI, Brahmer A, Ehlert T, Kluge K, Philippi KFA, Boedecker SC, et al. Validating quantitative PCR assays for cfDNA detection without DNA extraction in exercising SLE patients. Sci Rep. 2021;11:13581.

82. Erpenbeck L, Schön MP. Neutrophil extracellular traps: protagonists of cancer progression? Oncogene. 2017;36:2483–90.

83. Kos K, de Visser KE. Neutrophils create a fertile soil for metastasis. Cancer Cell. 2021;39:301–3.

84. Munir H, Jones JO, Janowitz T, Hoffmann M, Euler M, Martins CP, et al. Stromal-driven and Amyloid β-dependent induction of neutrophil extracellular traps modulates tumor growth. Nat Commun. 2021;12:683.

85. Nolan E, Malanchi I. Neutrophil “safety net” causes cancer cells to metastasize and proliferate. Nature. 2020;583:32–3.

86. Yang L, Liu Q, Zhang X, Liu X, Zhou B, Chen J, et al. DNA of neutrophil extracellular traps promotes cancer metastasis via CCDC25. Nature. Nature Publishing Group; 2020;583:133–8.

87. Barnes BJ, Adrover JM, Baxter-Stoltzfus A, Borczuk A, Cools-Lartigue J, Crawford JM, et al. Targeting potential drivers of COVID-19: Neutrophil extracellular traps. J Exp Med. 2020;217.

88. Zuo Y, Yalavarthi S, Shi H, Gockman K, Zuo M, Madison JA, et al. Neutrophil extracellular traps in COVID-19. JCI Insight. 2020;5.

